# Tuning the stability of DNA tetrahedra with base-stacking interactions

**DOI:** 10.1101/2024.06.10.598265

**Authors:** Jibin Abraham Punnoose, Dadrian Cole, Tristan Melfi, Vinod Morya, Bharath Raj Madhanagopal, Alan A. Chen, Sweta Vangaveti, Arun Richard Chandrasekaran, Ken Halvorsen

**Affiliations:** The RNA Institute, University at Albany, State University of New York, Albany, NY, USA; Department of Biological Sciences, University at Albany, State University of New York, Albany, NY, USA; Department of Chemistry, University at Albany, State University of New York, Albany, NY, USA; Department of Nanoscale Science and Engineering, University at Albany, State University of New York, Albany, NY, USA

**Author notes:** These authors contributed equally to this work (Jibin Abraham Punnoose, Dadrian Cole).

## Abstract

DNA nanotechnology relies on programmable anchoring of regions of single-stranded DNA through base pair hybridization to create nanoscale objects such as polyhedra, tubes, sheets, and other desired shapes. Recent work from our lab measured the energetics of base-stacking interactions and suggested that terminal stacking interactions between two adjacent strands could be an additional design parameter for DNA nanotechnology. Here, we explore that idea by creating DNA tetrahedra held together with sticky ends that contain identical base pairing interactions but different terminal stacking interactions. Testing all 16 possible combinations, we found that the melting temperature of DNA tetrahedra varied by up to 10 °C from altering a single base stack in the design. These results can inform stacking design to control DNA tetrahedra stability in a substantial and predictable way. To that end, we show that a 4 bp sticky end with weak terminal stacking does not form stable tetrahedra, while strengthening the stacks confers high stability with a 46.8 ± 1.2 °C melting temperature, comparable to a 6 bp sticky end with weak stacking (49.7 ± 2.9 °C). The results likely apply to other types of DNA nanostructures and suggest that terminal stacking interactions play an integral role in formation and stability of DNA nanostructures.

---

The advent of DNA nanotechnology has brought the opportunity to design and build materials from DNA, with nanoscale precision to build desired shapes with sizes ranging from nanometers to microns.^1–3^ The use of DNA and RNA as programmable materials can be attributed to the predictable base-pairing which defines interacting regions, as well as the stark (∼50 fold) difference in the stiffness between single and double stranded DNA.^4^ This allows a combination of structural rigidity in double stranded regions and tight bends in single-stranded regions. Structures can be formed by folding long single-stranded DNA using short DNA oligos called staples (in DNA origami) or by the self-assembly from smaller motifs through sticky end cohesion (hierarchical assembly).^5,6^ In both cases, hybridization of short DNA segments brings selected regions or other short fragments together in a desired geometry. Over the past three decades, the field has grown considerably, and hundreds of structures of varying complexity and function have been designed for applications including drug delivery, molecular computation, and nanorobotics.^7–9^

The core design principles in DNA nanotechnology involve hybridization and routing of various DNA strands between different regions throughout the structure. The contact areas between strands in individual uninterrupted duplex regions are typically short, ranging from a few base pairs to ∼20 or more base pairs depending on design and assembly technique.^5,6^ Complex designs including DNA origami can have several hundreds of such contacts, leaving a similar number of nicks and junctions that can potentially impact the structure and stability of the DNA nanostructures. Efforts to enhance the strength at these regions have typically included chemical (eg: crosslinking^10^) or enzymatic processes (eg: ligation^11^). One aspect that is rarely considered is how terminal base stacking interactions at nicks contribute to the overall stability of assembled DNA nanostructures.

Base stacking interactions have been considered and studied in DNA nanostructures, typically in the joining of individual structures into multimers. In the foundational DNA origami work, blunt end stacking was observed to uncontrollably join individual DNA origami structures. This was mostly viewed as problematic, and solved by passivating the ends or by omitting the staple strands on the edges of DNA origami structures.^5^ Since then, the effect of base stacking interactions on DNA nanostructure assembly has been noted to cause the formation of 1D arrays from blunt-ended 3-helix DNA motifs^12^ that otherwise form large, uniform arrays when containing sticky ends. Several surface-based assembly of 2D DNA origami lattices show the possibility of creating large-scale, ordered DNA arrays with stacking interactions rather than sticky ends.^13–15^ The significance of base stacking in DNA nanostructures was also demonstrated by a single molecule study that used DNA origami beams with an interface containing multiple base stacks in DNA bundles to quantify pairs of base stacking interactions.^16^

Partly inspired by that previous work^16^ and interested to further disentangle individual base stacking energies from dinucleotide pairs, we recently quantified individual base-stacking energies^17^ using a high-throughput single molecule technique called the centrifuge force microscope (CFM).^18,19^ We found energies of single nucleotide stacking interactions ranging from −0.5 to −2.3 kcal/mol, generally larger than predicted from the dinucleotide pair measurements^16,20^ but consistent with a single A|G stack measurement.^21^ The results suggested that terminal base-stacking energies are sufficiently large to influence the stability of short sticky ends, including those used in ligation and hierarchical assembly of DNA nanostructures.

Considering the recent information on single-nucleotide stacking energetics, and our still limited understanding of how interfacial base-stacks might influence the stability of individual DNA nanostructures, we set out to provide a more systematic study on if base stacking interactions might be used as a design parameter. In this paper, we provide a comprehensive examination of the role of base-stacking interactions on the stability of a model DNA nanostructure – the DNA tetrahedron. The DNA tetrahedron is among the most studied nanostructures for applications including drug delivery and sensing.^22^ While there are a few different designs,^6,23^ we chose the DNA tetrahedron which is hierarchically self-assembled from four identical 3-point-star motifs that connect to each other through pairs of sticky ends (Figure 1). In our previous work, we used a 3-point-star motif with 4 bp sticky ends for proof-of-concept demonstration that terminal base-stacking can impact stability.

**Figure 1.**
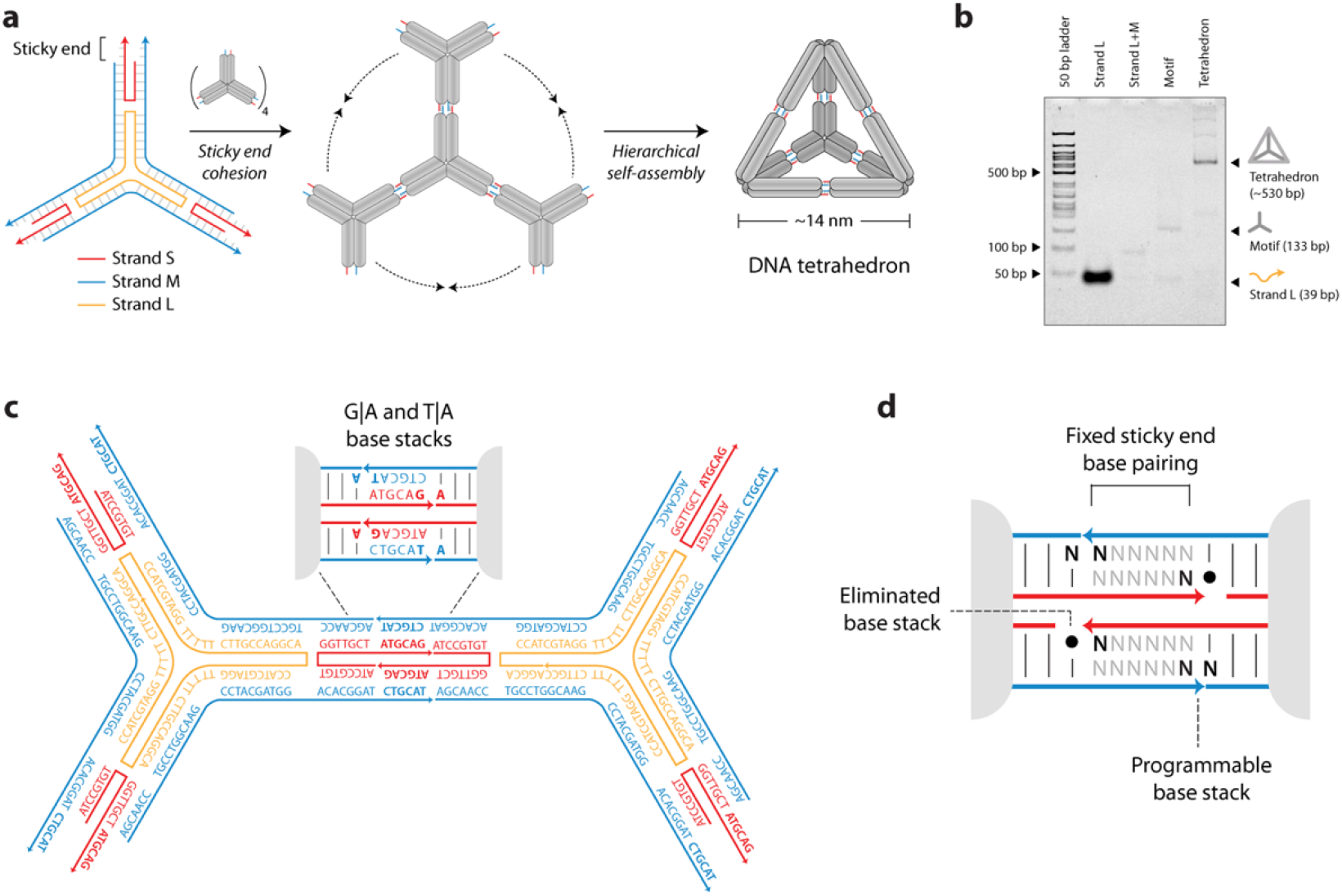
Self-assembly of 3-point-star motif into a DNA tetrahedron. (a)The scheme illustrates the hierarchical self-assembly of the 3-point-star motifs using sticky-end cohesion into a DNA tetrahedron. The typical tetrahedra consists of a pair of 4-nt sticky ends on each arm, which when assembled would each form four base-pairs and two base-stacks. (b) Formation of tetrahedron with 4-nt sticky ends confirmed using non denaturing PAGE. (c) Detailed design of a representative 3-point-star motif used in this study with 6-nt sticky ends and with a G|A and a T|A stack. (d) Detailed design of sticky-end indicating positions of programmable stacks and eliminated nucleotides to isolate an individual base-stack’s contribution to the overall stability.

To test the stability of DNA tetrahedra with all 16 combinations of terminal base stacking interactions, we first had to design sequences for the tetrahedra that could 1) preserve identical base-pairing interactions in the sticky ends across all variants, and 2) enable robust construction of the DNA tetrahedron for all variants. Our previous proof-of-concept work used a 4 bp sticky end (Figure 1a,b) but we found that 3-point-star motifs failed to assemble into tetrahedra with weaker base-stacks.^17^ To address this issue, we needed to strengthen the sticky ends of the 3 point star motifs. We tested 5 and 6 bp sticky-ends for a relative strong and weak base-stack and assessed the formation (Figure S1 & S2). From these experiments, we moved forward with a design incorporating 6 bp sticky ends (Figure 1c). To individually test each base stack, we omitted one terminal base stack from the design by shortening the 5’ end of one strand by 1 nt (outside of the sticky end region). We varied the other base stack by altering the downstream base outside the sticky end on either the short or medium strands (Figure 1d). This design allows a fair comparison between all base-stacks with unaltered sequence in the base pairing region of the sticky ends for all the 3-point-star motifs. This strategy provides a practical way to isolate the effect of changing an individual stacking interaction with minimal perturbation to the overall design.

We assembled the tetrahedra by mixing DNA strands L, M, and S (sequences in Table S1) in a 1:5:5 ratio at 30□nM in Tris-Acetic-EDTA-Mg^2+^ (TAE/Mg^2+^) buffer (40□mM Tris base (pH 8.0), 20□mM acetic acid, 2□mM EDTA, 12.5□mM magnesium acetate). The DNA solution was slowly cooled from 95□°C to room temperature over 48□h in a 2 liter water bath insulated in a Styrofoam box (Figure S2). All tetrahedra used the same L strand, but DNA tetrahedra with different base stacks had varying M and S strands to achieve the different stacking configurations (Figure 2a, Table S2). We validated the formation of each DNA tetrahedron for all base-stack combinations using non-denaturing 4% polyacrylamide gel electrophoresis (Figure 2b).

**Figure 2.**
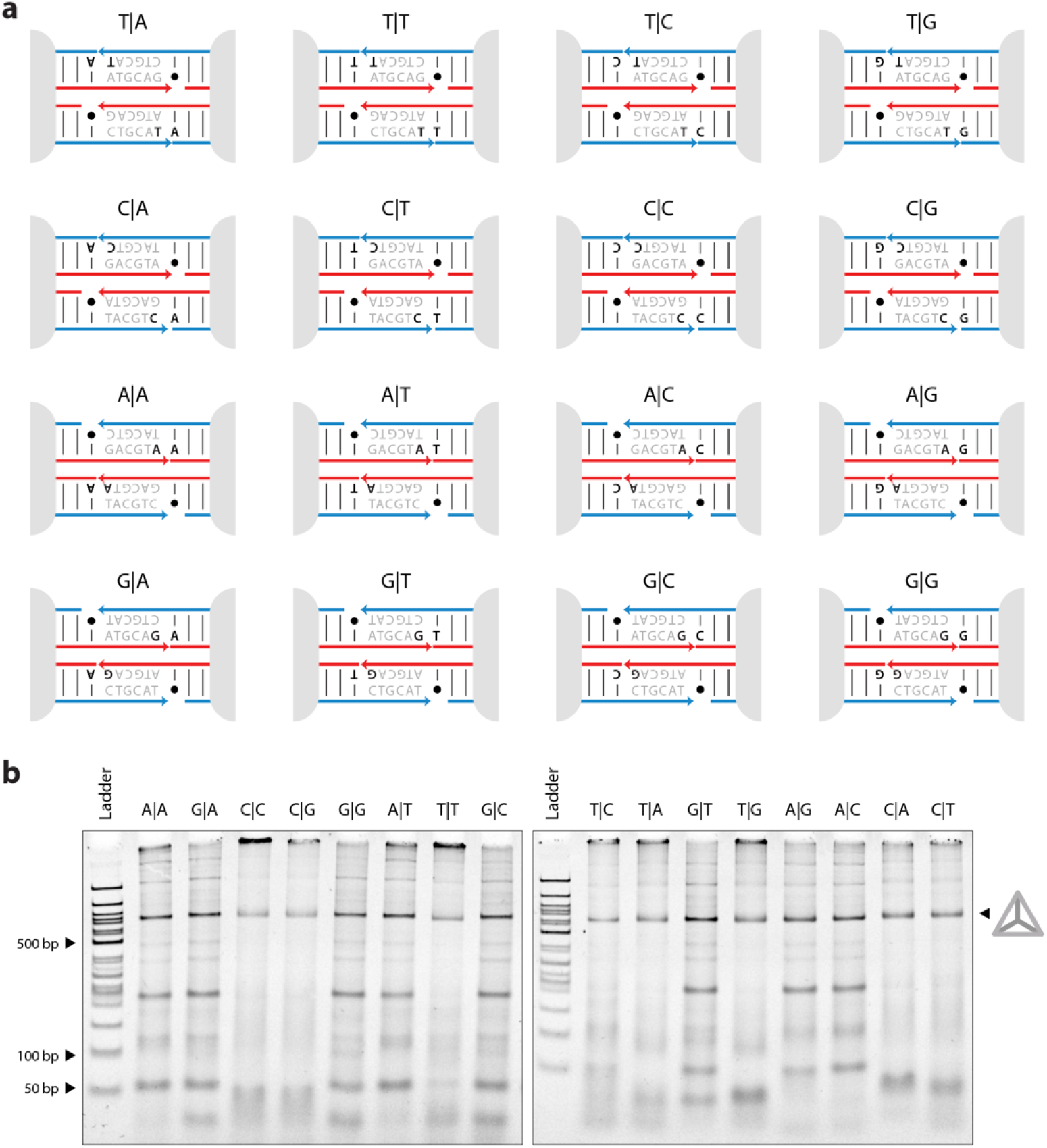
Design of strands and validation of DNA tetrahedra assembly. (a) Schematics of stick ends and strand designs for all the 16 tetrahedra with differing terminal stacking interactions. Stacked bases upon assembly are bolded and “•” represents a gap. (b) Formation of all 16 tetrahedra confirmed using non-denaturing PAGE.

While it can be tempting to draw conclusions based on the assembly of the tetrahedra, the efficiency in making the final product can be influenced by strand stoichiometry, strand purity, mixing precision, and hybridization kinetics. These can be difficult to accurately control or assess for 16 independent mixtures of L, M, and S strands drawn from 33 different oligo pools. We considered that a more reliable metric would be to measure the thermal stability of the tetrahedra that did form, independently normalized to the room temperature control for each variant. To assess this thermal stability, we incubated the formed structures for an hour in a range of temperatures (Figure 3a). Following thermal incubation, the fraction of tetrahedra remaining intact was determined by running the sample on a 4% non-denaturing PAGE (Figure 3b and S3-S18), and quantifying the band intensity at each temperature normalized against the intensity at room temperature. For each tetrahedra, the data followed a sigmoidal curve, with increased thermal dissociation observed at higher temperatures until no tetrahedra remained (Figure 3c).

**Figure 3.**
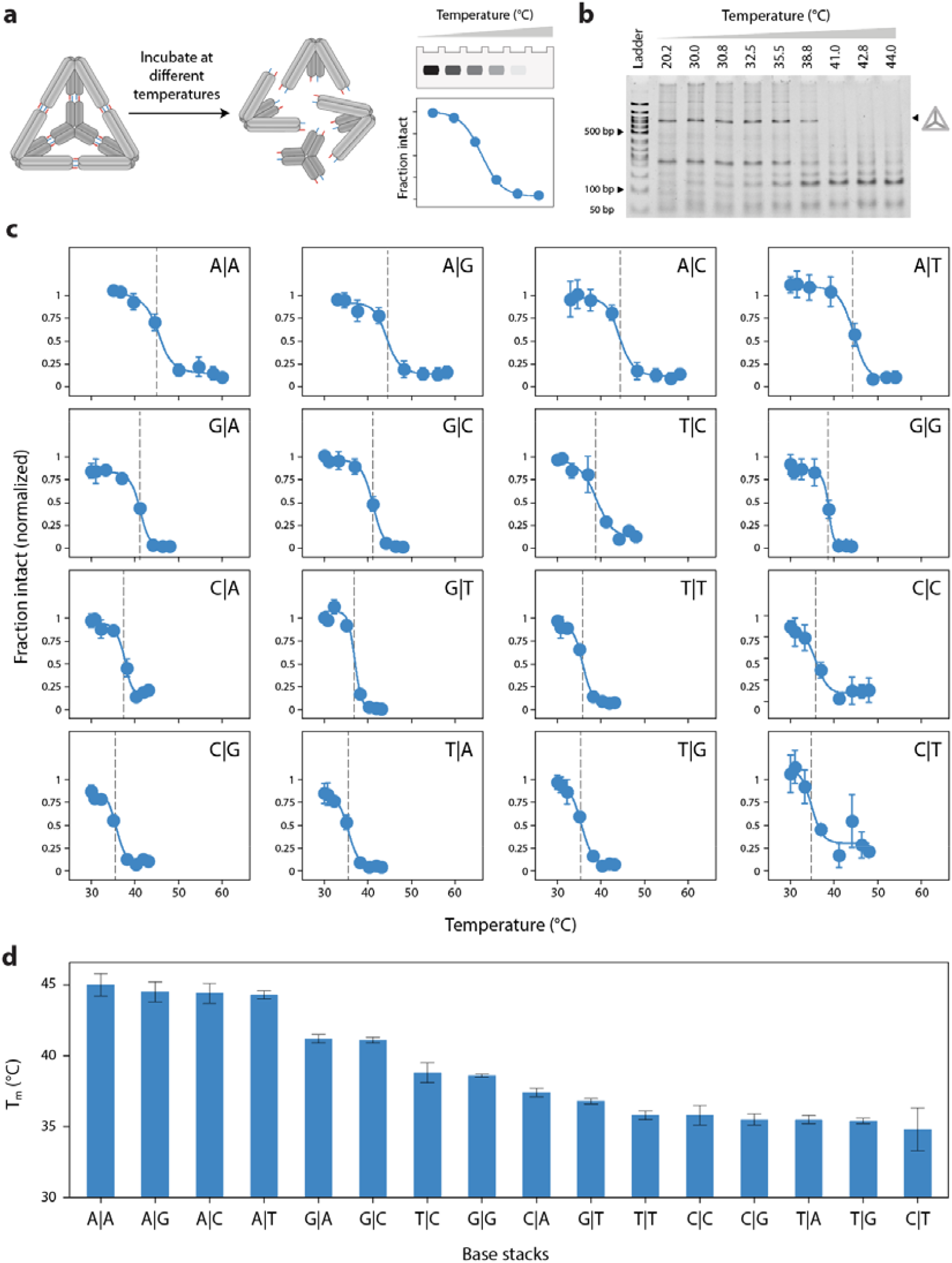
Thermal tolerance of DNA tetrahedra. (a) Cartoon illustration showing disassembly of DNA tetrahedron at elevated temperatures and subsequent analysis of the intact fraction using a non-denaturing PAGE. (b) Non-denaturing PAGE showing the change in intact fraction of tetrahedron with a G|G stack when incubated in a range on temperature from 20.2 °C to 44.0 °C for an hour. (c) The melting temperature is obtained by fitting a sigmoidal Boltzmann function to the intact fraction of DNA tetrahedron plotted against the incubation temperature. The fitting line represents the average of the individual fit from triplicate experiments (Figure S3-S18) and the T_m_ is the average T_m_ obtained from the three individual fits. (d) Bar graph representation of the average melting temperatures and the error bar represents the propagated error from the individual fits.

To compare the stability of the tetrahedra, we determined an apparent melting temperature by fitting each plot of intact fraction vs temperature with a Boltzmann function. Each of the 16 tetrahedra were independently prepared and analyzed in experimental triplicates to obtain mean melting temperatures (Figure 3c, Figure S2-S17). These results show a range of melting temperatures spanning ∼10 °C, ranging from 45.0 ± 0.8 °C for the A|A stack to 34.8 ± 1.5 °C for the C|T stack. In general, we observed that the most stable tetrahedra were those having a purine - purine stack while the least stable were those having a pyrimidine – pyrimidine stack (Figure 3d). These results generally followed similar trends as the energetics we found in our previous work quantifying base stacking energies,^17^ but do not match exactly. The differences between the rank order strengths of base-stacking interactions between this study and our previous may arise because of differences between thermal and force induced dissociation, as well as inherent difference in complexity between a nanostructure and a simple duplex.

One surprising result from this work was the apparent asymmetry between certain pairs of stacks with different polarities, for example A|T vs. T|A. In our previous work,^17^ we assumed that polarity would not affect the stacking energy but we did not explicitly test this assumption. To further explore this, we ran all-atom MD simulations of a nicked duplex with either an A|T or T|A stack at the interface at six different temperatures between 300 K and 400 K (Figure 4a-d). At lower temperatures (300 K and 320 K), we observed A|T and T|A distances at ∼0.25 nm throughout the simulation, though the T|A stack transiently sampled some longer distance configurations. However, at 340 K and higher the T|A stack falls apart as observed by distance between the bases greater than 0.5 nm for the majority of the simulation. Such behavior is observed for the A|T stack only at 400 K. These observations support our experimental evidence that the A|T configuration is more stable than the T|A configuration as the A|T stack remains intact at relatively higher temperatures (Figure 4e).

**Figure 4.**
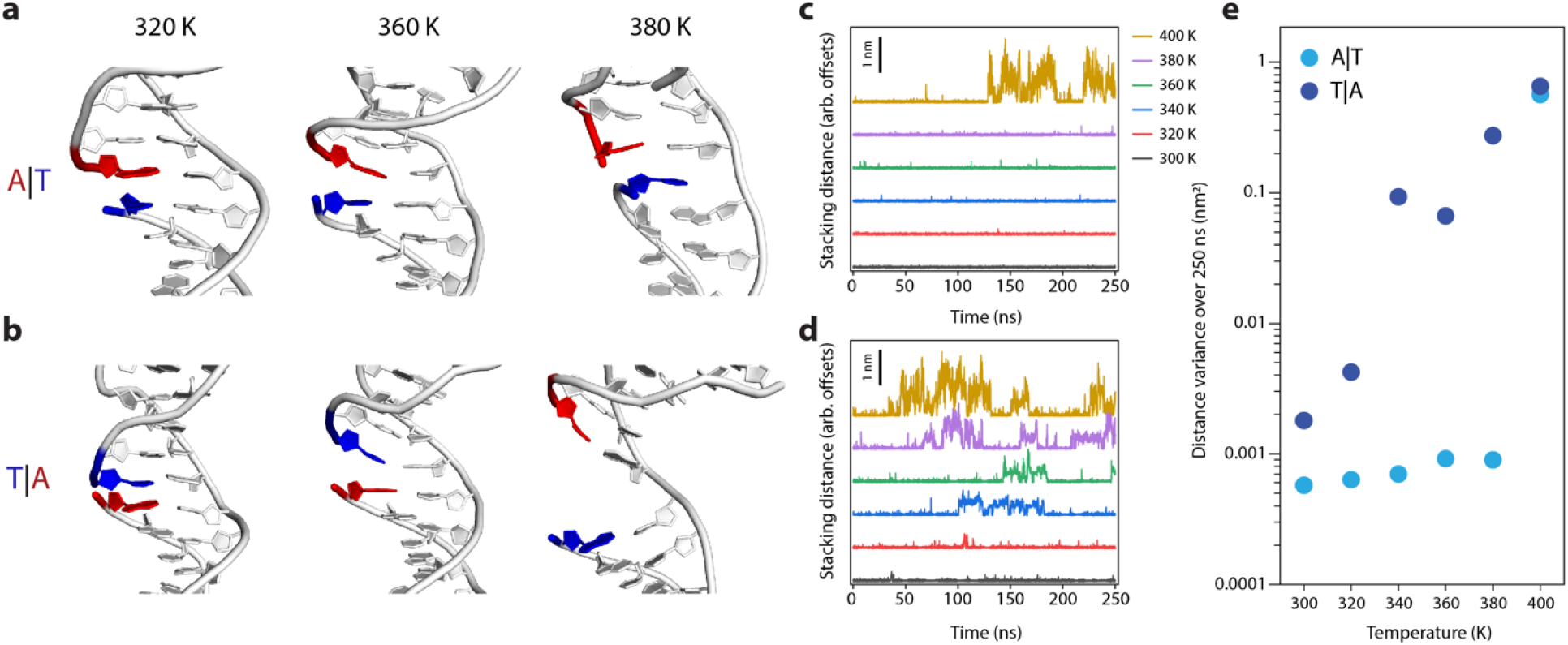
Molecular dynamics simulations of A|T vs. T|A stacking interfaces. (a) Representative snapshots from the simulation at various temperatures for A|T, and (b) for T|A. (c) time dependent stacking distances for A|T, and (d) for T|A. (e) Variance in the stacked distance over the duration of the simulation for each condition.

Our results indicate that in the context of a DNA tetrahedron, changing a single terminal base stacking interaction in a 6 bp sticky end can alter the melting temperature of the structure by as much as 10 °C. This clearly illustrates the importance of base-stacking in DNA nanostructure stability. We expect that the relative contribution of base-stacking interactions will increase as the length of the sticky ends get shorter, and also be more pronounced when both terminal stacking interactions are present in the sticky ends (as opposed to the single interaction tested here).

To further test these ideas, we designed three additional 3-point-star motifs with sticky ends that fully base stack on both sides with their neighboring strands. We designed two 4 bp sticky ends with either weak stacking (T|T + T|T) or strong stacking (A|A + A|A) and another 6 bp sticky end with weak stacking (G|T + T|T) (Figure 5a). We found that the tetrahedra with 6 bp sticky ends and weak base-stacks and 4 bp with strong base stacks had similar melting temperatures of 49.7 ± 2.9 °C and 46.8 ± 1.2 °C, respectively, while the 4 bp sticky ends with weak base stacks did not form stable tetrahedra (Figure 5b-c & S22. It is worth comparing these against the 6 bp tetrahedra with a single G|T or T|T stack, (melting temperatures of 36.8 ± 0.2 ° C and 35.8 ± 0.3 °C, respectively), which were considerably less stable than either 6 bp with both G|T and T|T base-stacks or a 4 bp with a pair of stronger A|A base-stacks. We tested a few additional configurations as well (Figure S21 & S23). These experiments indicate that strong base-stacks can largely compensate for the reduced binding energy from a shortened sticky-end, at least in the context of these tetrahedra.

**Figure 5.**
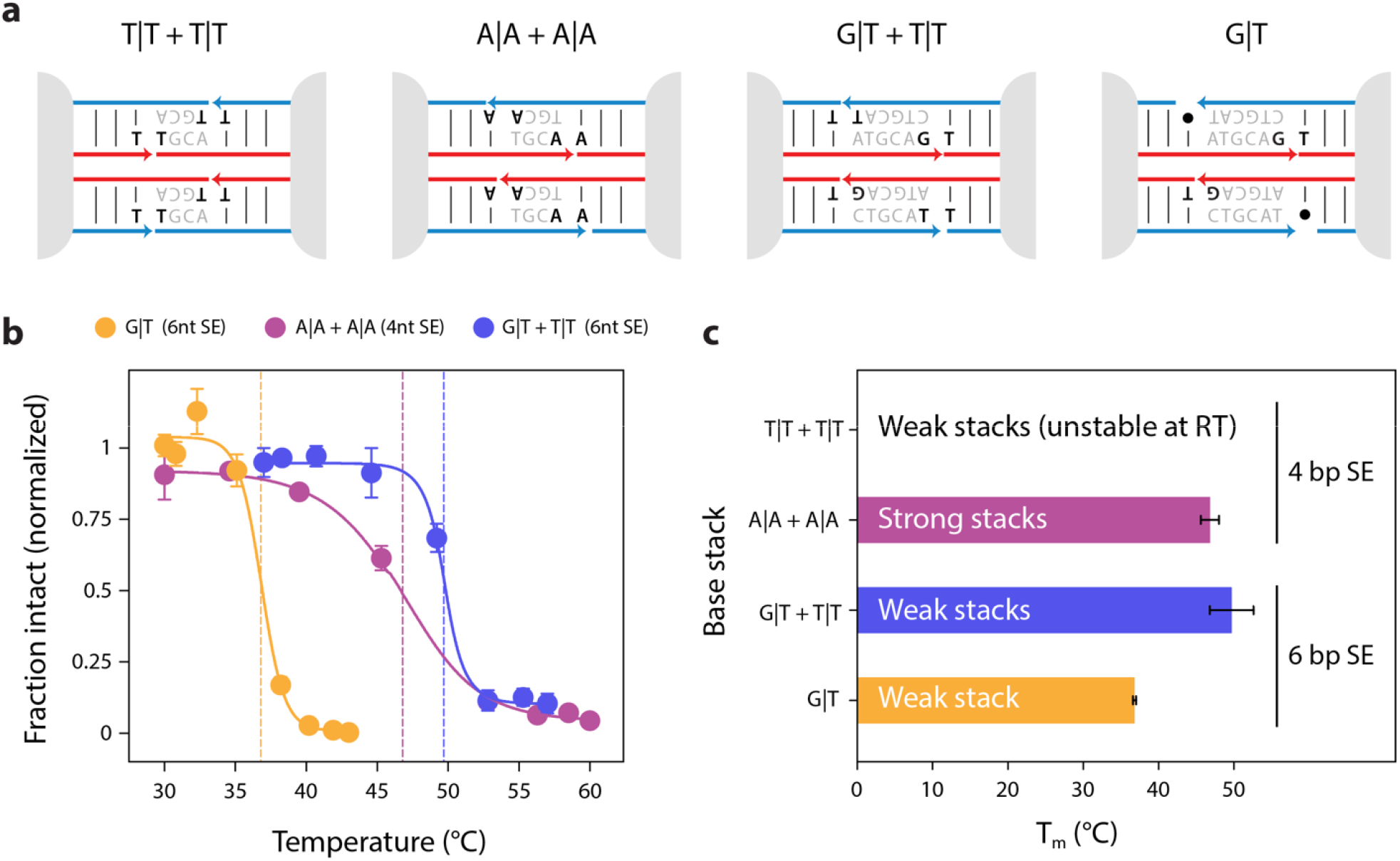
Tuning the thermal stability of DNA tetrahedra. (a) Schematics of sticky ends and strand designs for tetrahedra with 4 bp sticky ends with weak or strong stacks, 6 bp sticky ends with weak stacks, and with one stack omitted (from Figure 2) for comparison. (b) The average melting profiles of the 3 tetrahedra that formed from triplicate measurements (Figures S18, S20-S22), fit with a Boltzmann curve to determine the melting temperatures. (c) A bar graph representation of the average melting temperatures, with error bars representing the propagated error from the individual fits.

As we have shown, base-stacks formed by sticky-end cohesion can affect the stability of DNA tetrahedra. Our work further elucidates some key principles for sticky ends in DNA tetrahedra: 1) Generally, purine-purine stacks, especially when adenine is on the 5’ site yield the most thermally stable tetrahedra (and conversely less stable for pyrimidine-pyrimidine stacks); 2) The omission of one stack in the design (while maintaining the same-base pairs) can greatly reduce the stability of DNA tetrahedra; and 3) Stronger base stacking can partly compensate for reduced stability when shortening sticky ends.

Our study has some limitations which are also worth discussing. We used unpurified tetrahedra which may contain byproducts including some aggregates and partly formed structures. We argue that these byproducts should not interfere with our analysis due to their different size distributions that separate them in the gels. Gel-based purification would introduce new uncertainties including 1) potential changes in stability due to dye incorporation and UV light damage, 2) potential contamination with nucleases or changes in buffer composition, and 3) potential variations in final concentrations (and thus melting temperatures) due to purification yields. Analysis by gel provides a good way to quantify the overall amount of intact tetrahedra without requiring purification. Other methods may provide real-time analysis or more detailed information about disassociation pathways, but also come with tradeoffs. Fluorophore-quencher pairs could in principle provide real time data of well-purified structures, but fluorophores may affect the very interactions being measured,^17,24^ and the relationship between overall structure and the binding status of individual strands (or portions of strands) is not always obvious. For example, it may be possible for a tetrahedron to remain largely intact even with a single strand unbound and conversely a disassembled tetrahedron does not guarantee that all strands are dehybridized. Other methods may be considered for such measurements but can prove difficult for such small structures including light scattering techniques^25,26^ and AFM imaging.

Overall, our work suggests that base stacking interactions can be part of the design considerations for some types of DNA nanostructures. DNA origami is likely to be the least affected due to typically larger duplex regions, but hierarchical assembly,^6,27^ blunt end assembly,^12,28^ DNA bricks,^29^ DNA superstructures,^30^ and DNA lattices^31^ are all likely to be affected by base stacking interactions at interfaces. As we demonstrated here for the tetrahedron, it is possible to tune the stability of DNA nanostructures by altering key stacking interactions. This may offer new strategies for strengthening (or weakening) structures, in addition to other current tools including ligation ^32^ and modified DNAs.^11,33^

## Acknowledgements

The authors acknowledge general support from The RNA Institute, and to the University at Albany Summer Research Program (UASRP) for support for D.C. Research reported in this publication was supported by the National Institutes of Health (NIH) through the National Institute of General Medical Sciences (NIGMS) under awards R35GM124720 to K.H (and supplement-08S1 to D.C.) R35GM150672 to A.R.C. and R35GM133469 to A.A.C. The content is solely the responsibility of the authors and does not necessarily represent the official views of the NIH.

## Author contributions

The project was conceived and planned by J.A.P. and K.H. with assistance from A.R.C. Strands were designed by J.A.P. and A.R.C. Experiments were designed by J.A.P., A.R.C. and K.H. Experiments were performed by D.C., V.M., B.R.M., and supervised by J.A.P. Data was analyzed by J.A.P, D.C, and V.M. Molecular dynamics simulations were performed and analyzed by T.M., S.V., and A.C. The project was supervised by K.H. Figures were made by J.A.P. and A.R.C. The paper was written by J.A.P., A.R.C., and K.H. with general input and editing from all authors.

