## Supplementary Information for "Tuning the stability of DNA tetrahedra with base-stacking interactions"

### Table of Contents:

Supplementary Methods: Tetrahedron assembly, melting and analysis, MD
Table S1: List of oligonucleotides
Table S2: Combination of oligonucleotides to form the various tetrahedra
Figure S1: Gel images of tetrahedrons with 5nt and 6nt sticky end
Figure S2: Annealing protocol for DNA tetrahedra.
Figure S3: Gel images of tetrahedron with A|A base-stack at various temperatures
Figure S4: Gel images of tetrahedron with A|C base-stack at various temperatures
Figure S5: Gel images of tetrahedron with A|G base-stack at various temperatures
Figure S6: Gel images of tetrahedron with A|T base-stack at various temperatures
Figure S7: Gel images of tetrahedron with C|A base-stack at various temperatures
Figure S8: Gel images of tetrahedron with C|C base-stack at various temperatures
Figure S9: Gel images of tetrahedron with C|G base-stack at various temperatures
Figure S10: Gel images of tetrahedron with C|T base-stack at various temperatures
Figure S11: Gel images of tetrahedron with G|A base-stack at various temperatures
Figure S12: Gel images of tetrahedron with G|C base-stack at various temperatures
Figure S13: Gel images of tetrahedron with G|G base-stack at various temperatures
Figure S14: Gel images of tetrahedron with G|T base-stack at various temperatures
Figure S15: Gel images of tetrahedron with T|A base-stack at various temperatures
Figure S16: Gel images of tetrahedron with T|C base-stack at various temperatures
Figure S17: Gel images of tetrahedron with T|G base-stack at various temperatures
Figure S18: Gel images of tetrahedron with T|T base-stack at various temperatures
Figure S19: Gel images of tetrahedron with G|T and T|T base-stack at various temperatures
Figure S20: Gel images of tetrahedron with 2 A|A base-stack & 4-nt sticky end at various temperatures
Figure S21: Triplicate melting experiments with tetrahedron with 1 T|T stack and a 5 nucleotide sticky end
Figure S22: Gel images showing formation of different 6 and 4-nt tetrahedron with various base-stacks
Figure S23: Thermal stability difference between tetrahedron with T|T stacks and A|A stacks.

### Supplementary Methods

#### *Tetrahedra construction*

To assemble the DNA tetrahedron, desalted L, M and S strands (Table S1) were mixed in a 1:5:5 ratio at a final concentration of 30nM in Tris-acetate-EDTA-$Mg^{2+}$ buffer (40mM Tris base, pH 8.0; 20mM acetic acid; and 12.5mM magnesium acetate). The DNA samples were then annealed by placing them in a 2L water bath within a Styrofoam box, which allowed for their gradual cooling from 90°C to room temperature over a 48-hour period. To assemble DNA tetrahedra with different base stacks varying strand combinations were used (Table S2).

For one set of experiments, a thermal cycler was used to assemble the tetrahedra. To program the thermal cycler, we first measured the cooling rate of the thermal bath in Styrofoam using a thermometer (Figure S2). The thermal cycler was then programmed with the following cooling profile: 85 °C to 75 °C at a rate of -1 °C per 9 minutes, 75 °C to 60 °C at a rate of -1 °C per 12 minutes, 60 °C to 51 °C at a rate of -1 °C per 20 minutes, 51 °C to 43 °C at a rate of -1 °C per 20 minutes, 43 °C to 38 °C at a rate of -1 °C per 36 minutes, 38 °C to 34 °C at a rate of -1 °C per 45 minutes, 34 °C to 32 °C at a rate of -1 °C per 90 minutes, 32 °C to 31 °C at a rate of -1 °C per 180 minutes, 31 °C to 30 °C at a rate of -1 °C per 180 minutes, and 30 °C to 23 °C at a rate of -1 °C per 200 minutes.

#### *Thermal analysis*

To analyze the thermal stability of DNA tetrahedra, 5 µL of each tetrahedron was placed in nine tubes and incubated in a T100 Thermal Cycler (Bio-Rad) for one hour at predetermined experimental temperatures. After incubation, 5 µL of the samples were mixed with 1 µL of a glycerol based gel loading dye, and 5 µL total was used for each sample. Samples were run at 4 °C on 4% non-denaturing polyacrylamide (29:1 acrylamide/bisacrylamide) gels at 100V for one hour. Following electrophoresis, gels were stained in 50 mL of 1x Tris-acetate-EDTA-$Mg^{2+}$ buffer and 5 µL of 10,000X GelRed (Biotium) before imaging. Imaging was done with an Azure Biosystems Imager using the default settings for Ethidium Bromide ultraviolet illumination.

The intact fraction of tetrahedra in the polyacrylamide gel was measured using the Bio-Rad ImageLab Software. The bands corresponding to the DNA tetrahedron with varied base stacks at different temperatures were quantified and normalized against the band at room temperature. These normalized values were plotted in OriginLab and the data was fitted using a Boltzmann function, $f(x) = (A2 - A1)\ (1 + e(x - x_0)/dx) + A1$ , where A1 is the lower asymptote, A2 is the upper asymptote, x is the independent variable, $x_0$ is the inflection point, and d*x is* a parameter associated with the width of the sigmoidal curve. The thermal stability was determined by using individual $x_0$ values which is typically considered to represent the midpoint of the transition from one state to another, commonly known as the melting point. Fitting uncertainty from each replicate was used to determine the error represented as the uncertainty of the average melting temperatures from three individual measurements.

### ***MD simulations***

To investigate the difference in stability between the 2 stacking configurations we simulated 2 nicked duplexes: one with A|T stacking and the other with T|A stacking at the nicked interface in all atom molecular dynamics simulations. After modelling the two duplexes in Molecular Operating Environment (MOE), they were then simulated using GROMACS 2019.4 (1) for 250 ns at temperatures between 300 K to 400 K in 20 K increments, performing a total of simulations. Each system was neutralized by adding $K^+$ and $Cl^-$ ions. The tumuc1 force-field (2) was used with the accompanying ion parameters for this system.

All of the molecular dynamics simulations incorporated the leap-frog algorithm to integrate the equations of motion. The systems used the velocity rescaling thermostat to maintain the temperature (3). Pressure was maintained at 1 atm using Parrinello-Rahman barostat (4) for the equilibration. Long-range electrostatic interactions were calculated using particle mesh Ewald (PME) algorithm with a cut-off of 1.0 nm (5) Lennard-Jones interactions were also truncated at 1.0 nm. Water molecules were represented using the TIP3P model (6), and the LINCS algorithm was used to constrain the motion of hydrogen atoms bonded to heavy atoms (7). Each system was subjected to energy minimization to prevent any overlap of atoms, followed by 100 ps temperature and pressure equilibration, leading to the final production run of 250 ns. The simulations were visualized using PyMOL and the MolecularNodes Blender plugin. The rmsd and distance tools from GROMACS (1) were used for analysis. The distance between the bases was defined as the distance between the center of mass of the heavy atoms on the nucleobases. Nucleotides were considered stacked if this distance was less than 0.3 nm.

### Table S1: List of oligonucleotides

| Name | Sequence (5'-3') | Length |
|---|---|---|
| Tet L | AGG CAC CAT CGT AGG TTT TTC TTG CCA GGC ACC ATC GTA GGT TTT TCT TGC CAG GCA CCA TCG TAG GTT TTT CTT GCC | 78 |
| TET-M-6SE TG | GGC AAC CTG CCT GGC AAG CCT ACG ATG GAC ACG GAT CTG CAT | 42 |
| TET-S-6SE TG | TCC GTG TGG TGG CCA TGC AG | 20 |
| TET-M-6SE TC | CGC AAC CTG CCT GGC AAG CCT ACG ATG GAC ACG GAT CTG CAT | 42 |
| TET-S-6SE TC | TCC GTG TGG TGG CGA TGC AG | 20 |
| TET-M-6SE TT | TGC AAC CTG CCT GGC AAG CCT ACG ATG GAC ACG GAT CTG CAT | 42 |
| TET-S-6SE TT | TCC GTG TGG TTG CAA TGC AG | 20 |
| TET-M-6SE TA | AGC AAC CTG CCT GGC AAG CCT ACG ATG GAC ACG GAT CTG CAT | 42 |
| TET-S-6SE TA | TCC GTG TGG TTG CTA TGC AG | 20 |
| TET-M-6SE AG | GCA ACC TGC CTG GCA AGC CTA CGA TGG ACA CGG ACT ACG TC | 41 |
| TET-S-6SE AG | GTC CGT GTG GTT GCA GAC GTA | 21 |
| TET-M-6SE AC | GCA ACC TGC CTG GCA AGC CTA CGA TGG ACA CGG AGT ACG TC | 41 |
| TET-S-6SE AC | CTC CGT GTG GTT GCA GAC GTA | 21 |
| TET-M-6SE CG | GGC AAC CTG CCT GGC AAG CCT ACG ATG GAC ACG GAT TAC GTC | 42 |
| TET-S-6SE CG | TCC GTG TGG TTG CCG ACG TA | 20 |
| TET-M-6SE GG | GCA ACC TGC CTG GCA AGC CTA CGA TGG ACA CGG ACC TGC AT | 41 |
| TET-S-6SE GG | GTC CGT GTG GTT GCA ATG CAG | 21 |
| TET-M-6SE AT | GCA ACC TGC CTG GCA AGC CTA CGA TGG ACA CGG AAT ACG TC | 41 |
| TET-S-6SE AT | TTC CGT GTG GTT GCA GAC GTA | 21 |
| TET-M-6SE CC | CGC AAC CTG CCT GGC AAG CCT ACG ATG GAC ACG GAT TAC GTC | 42 |
| TET-S-6SE CC | TCC GTG TGG TTG CGG ACG TA | 20 |
| TET-M-6SE GC | GCA ACC TGC CTG GCA AGC CTA CGA TGG ACA CGG AGC TGC AT | 41 |
| TET-S-6SE GC | CTC CGT GTG GTT GCA ATG CAG | 21 |
| TET-M-6SE AA | GCA ACC TGC CTG GCA AGC CTA CGA TGG ACA CGG ATT ACG TC | 41 |
| TET-S-6SE AA | ATC CGT GTG GTT GCA GAC GTA | 21 |
| TET-M-6SE CA | AGC AAC CTG CCT GGC AAG CCT ACG ATG GAC ACG GAT TAC GTC | 42 |
| TET-S-6SE CA | TCC GTG TGG TTG CTG ACG TA | 20 |
| TET-M-6SE GT | GCA ACC TGC CTG GCA AGC CTA CGA TGG ACA CGG AAC TGC AT | 41 |
| TET-S-6SE GT | TTC CGT GTG GTT GCA ATG CAG | 21 |
| TET-M-6SE GA | GCA ACC TGC CTG GCA AGC CTA CGA TGG ACA CGG ATC TGC AT | 41 |
| TET-S-6SE GA | ATC CGT GTG GTT GCA ATG CAG | 21 |
| TET-M-6SE CT | TGC AAC CTG CCT GGC AAG CCT ACG ATG GAC ACG GAT TAC GTC | 42 |
| TET-S-6SE CT | TCC GTG TGG TTG CAG ACG TA | 20 |
| TET-M-5SE CT | TAG CAA CCT GCC TGG CAA GCC TAC GAT GGA CAC GGA TTC GTC | 42 |
| TET-S-5SE CT | TCC GTG TGG TTG CTA GAC GA | 20 |
| TET-M-5SE GA | AGC AAC CTG CCT GGC AAG CCT ACG ATG GAC ACG GAT CTG CT | 41 |
| TET-S-5SE GA | ATC CGT GTG GTT GCT AAG CAG | 21 |
| TET-M-6SE GT+TT | TGC AAC CTG CCT GGC AAG CCT ACG ATG GAC ACG GAA CTG CAT | 42 |
| TET-M-5SE-2TT | TCGGA AGCAACC TGCCTGGCAAG CCTAC GATGG ACACGGTAT | 42 |
| TET-S-5SE-1TT | TCCGA ATACCGTGT GGTTGCT | 21 |

| | | |
|---|---|---|
| TET-M-4SE 2AA | AAG CAA CCT GCC TGG CAA GCC TAC GAT GGA CAC GGT ATT CGA | 42 |
| TET-S-4SE 2AA | ATA CCG TGT GGT TGC TTT CGA | 21 |
| TET-M-4SE 2TT | TCG AAA GCA ACC TGC CTG GCA AGC CTA CGA TGG ACA CGG TAT | 42 |
| TET-S-4SE 2TT | TCG AAT ACC GTG TGG TTG CTT | 21 |
| TET-M-0SE-2TT | AAGCAACC TGCCTGGCAAG CCTAC GATGG ACACGGTAT | 38 |
| TET-S-0SE-2TT | ATACCGTGT GGTTGCTT | 17 |

### Table S2: Combination of oligonucleotides to form the tetrahedron with a specific base-stack

| Tetrahedron (base-stack) | Sticky end Length | L Strand | M Strand | S Strand |
|---|---|---|---|---|
| TG | 6 nt | Tet L | TET-M-6SE TG | TET-S-6SE TG |
| TC | 6 nt | Tet L | TET-M-6SE TC | TET-S-6SE TC |
| TT | 6 nt | Tet L | TET-M-6SE TT | TET-S-6SE TT |
| TA | 6 nt | Tet L | TET-M-6SE TA | TET-S-6SE TA |
| AG | 6 nt | Tet L | TET-M-6SE AG | TET-S-6SE AG |
| AC | 6 nt | Tet L | TET-M-6SE AC | TET-S-6SE AC |
| CG | 6 nt | Tet L | TET-M-6SE CG | TET-S-6SE CG |
| GG | 6 nt | Tet L | TET-M-6SE GG | TET-S-6SE GG |
| AT | 6 nt | Tet L | TET-M-6SE AT | TET-S-6SE AT |
| CC | 6 nt | Tet L | TET-M-6SE CC | TET-S-6SE CC |
| GC | 6 nt | Tet L | TET-M-6SE GC | TET-S-6SE GC |
| AA | 6 nt | Tet L | TET-M-6SE AA | TET-S-6SE AA |
| CA | 6 nt | Tet L | TET-M-6SE CA | TET-S-6SE CA |
| GT | 6 nt | Tet L | TET-M-6SE GT | TET-S-6SE GT |
| GA | 6 nt | Tet L | TET-M-6SE GA | TET-S-6SE GA |
| CT | 6 nt | Tet L | TET-M-6SE CT | TET-S-6SE CT |
| CT | 5 nt | Tet L | TET-M-5SE CT | TET-S-5SE CT |
| AG | 5 nt | Tet L | TET-M-5SE CT | TET-S-5SE CT |
| GT + TT | 6 nt | Tet L | TET-M-6SE GT+TT | TET-S-6SE GT |
| AA + AA | 4 nt | Tet L | TET-M-4SE 2AA | TET-S-4SE 2AA |
| T\|T | 5 nt | Tet L | TET-M-5SE-2TT | Tet-S-5SE-1TT |
| T\|T + TT | 4 nt | Tet L | TET-M-4SE 2TT | TET-S-4SE 2TT |
| Motif | 0 | Tet L | TET-M-0SE-2TT | TET-S-0SE-2TT |

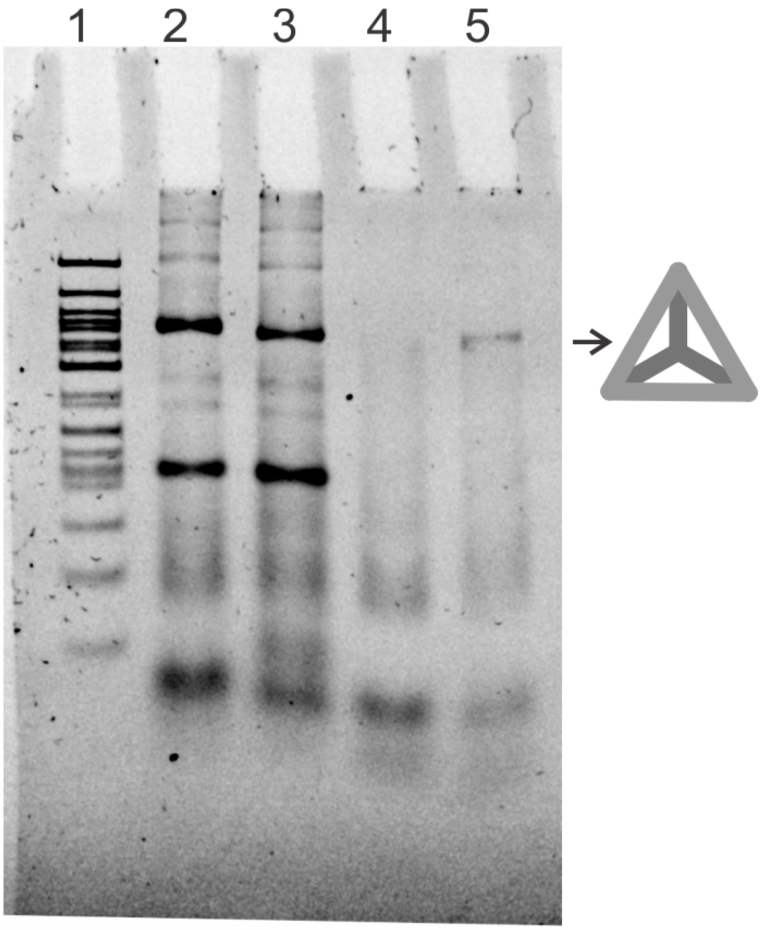


**Figure S1: Tetrahedron with 5 and 6 nt sticky ends.** Lane 1: 100 bp ladder, Lane 2: Tetrahedron with A|G base-stack and 5 nt sticky end, Lane 3: Tetrahedron with A|G base-stack and 6 nt sticky end, Lane 4: Tetrahedron with C|T base-stack and 5 nt sticky end, and Lane 5: Tetrahedron with C|T base-stack and 6 nt sticky end.

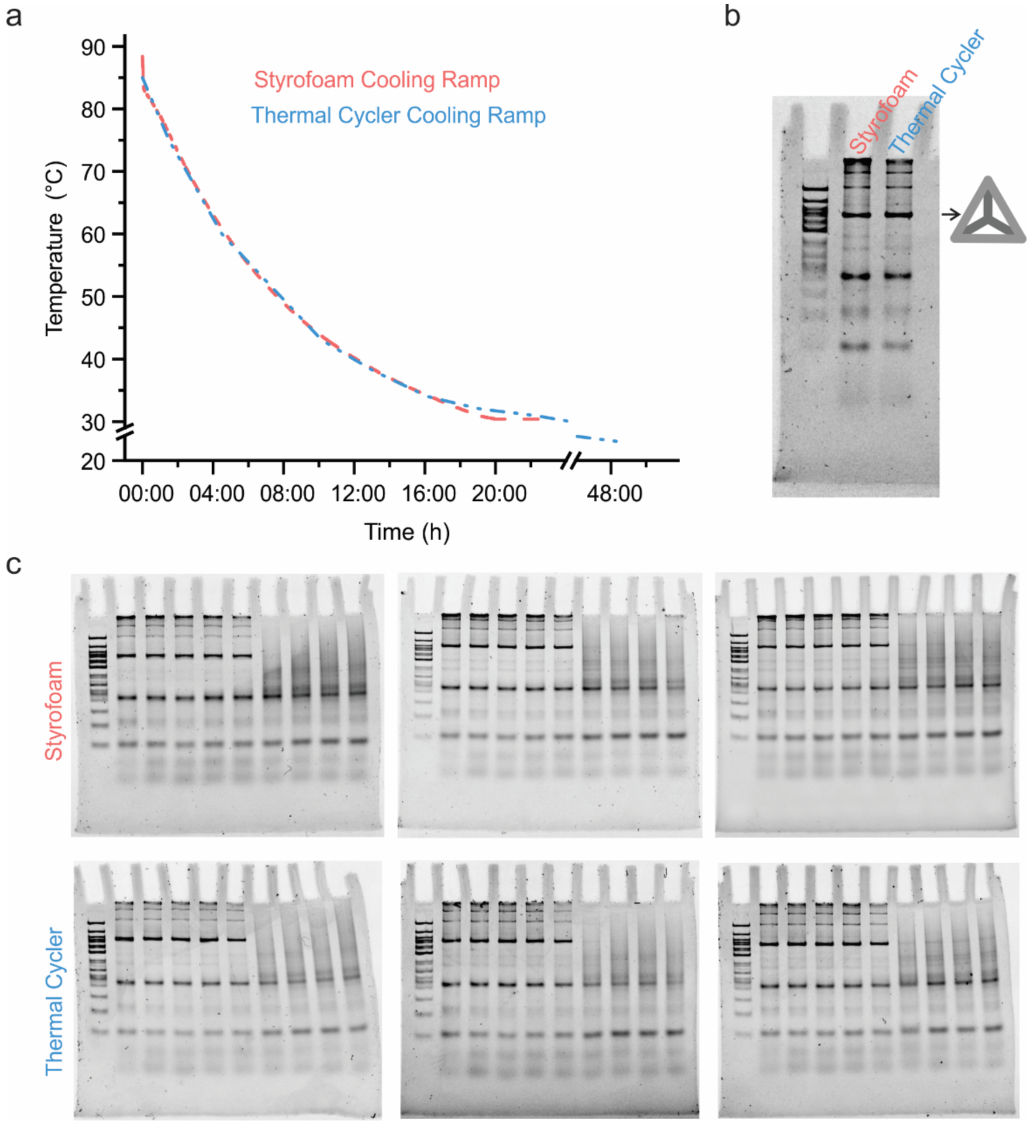


**Figure S2: Annealing protocol for DNA tetrahedra.** (a) Measured cooling rate of the thermal bath inside the Styrofoam box used for annealing DNA tetrahedra and the thermal cycler cooling profile developed to mimic the process. (b) Comparison of DNA tetrahedron formed using annealing in a Styrofoam and a thermal cycler. (c) Tetrahedron melting comparison using tetrahedron formed using Styrofoam box and a thermal cycler.

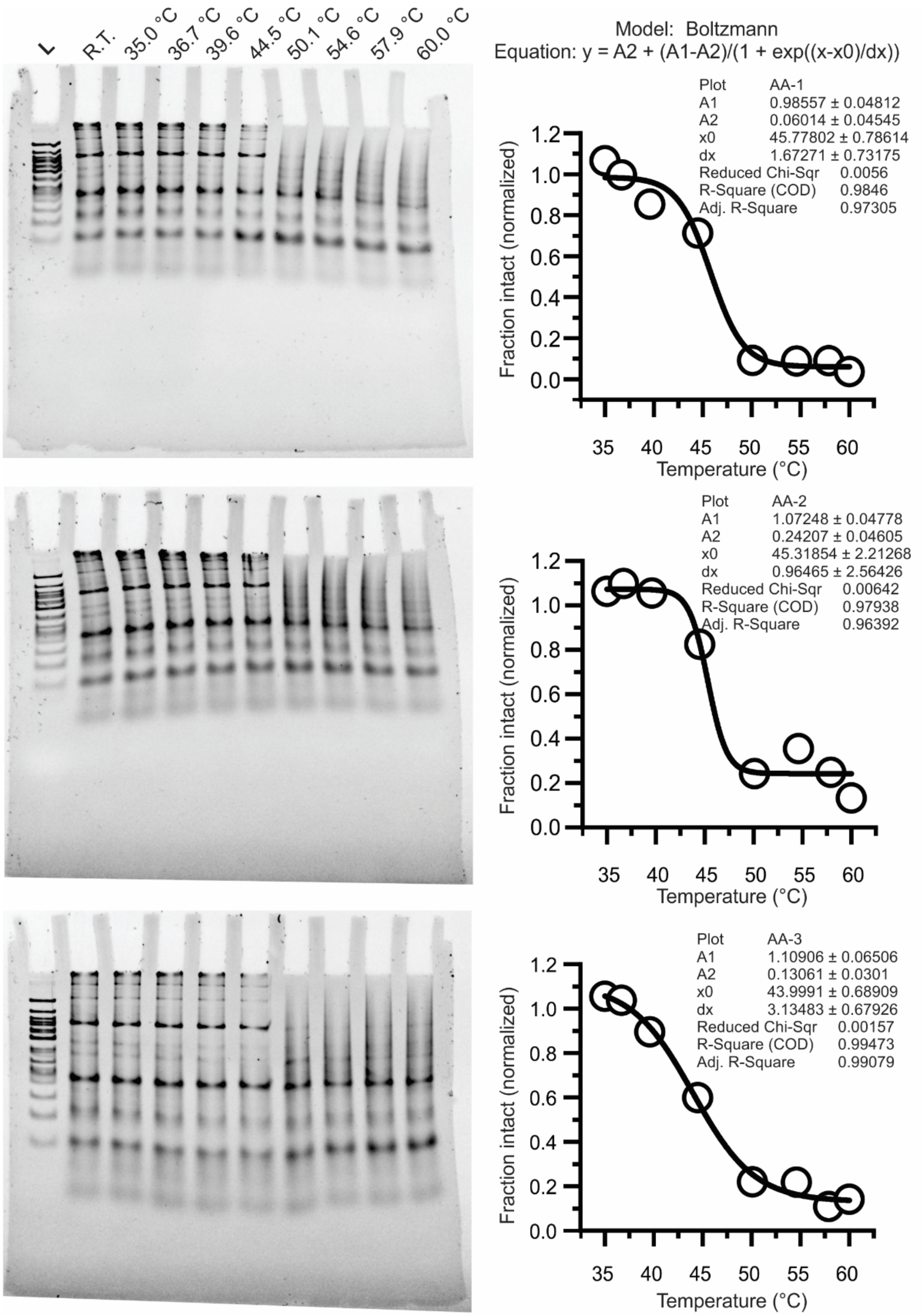


**Figure S3: Triplicate melting experiments with tetrahedron with an A|A stack with 6 nucleotide sticky end.** The formed nanostructures were incubated at various temperatures for 1 hour as indicated in the lane. The intact fraction of tetrahedron was fitted with Boltzmann model to obtain the melting temperature for each experiment.

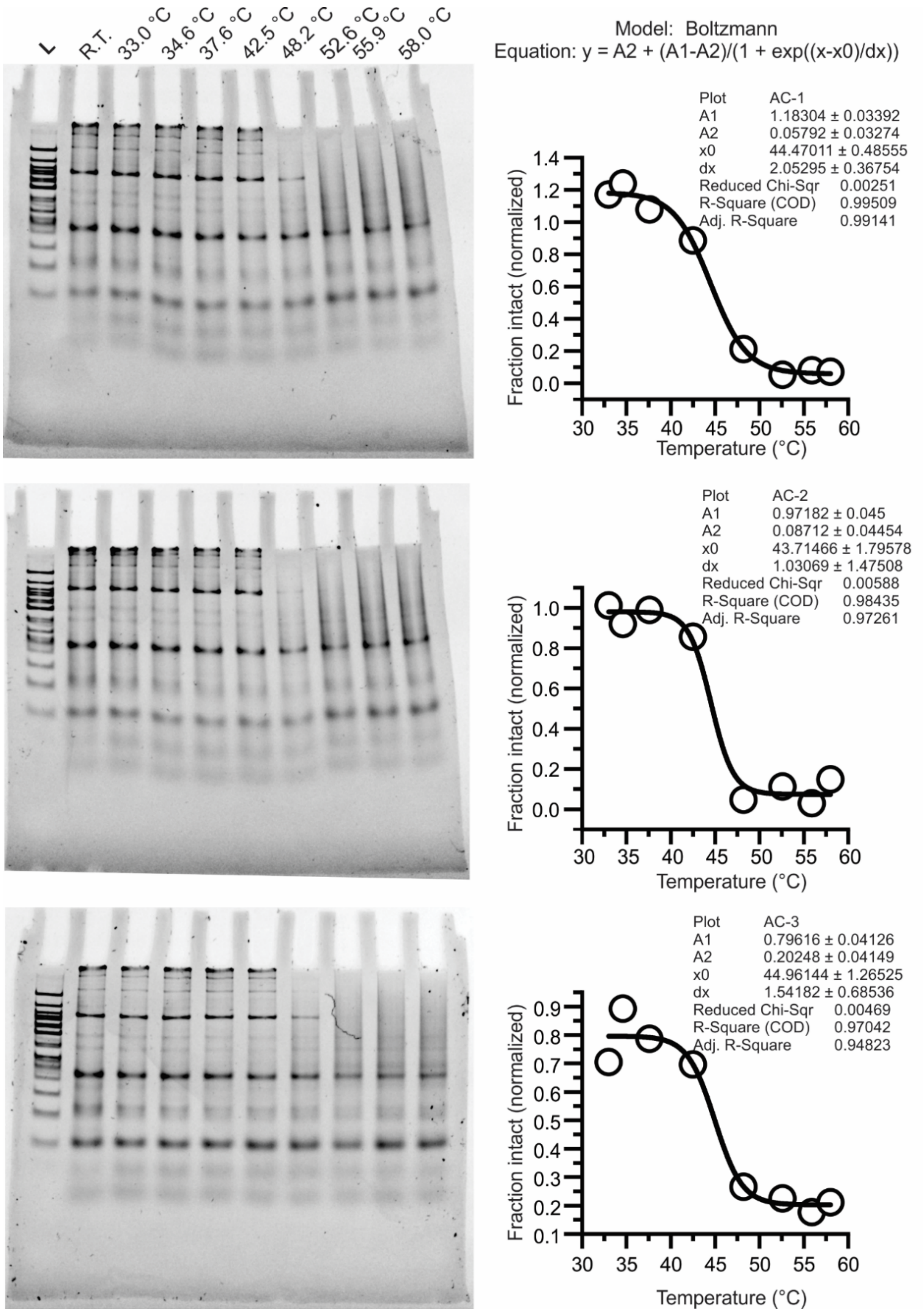


**Figure S4: Triplicate melting experiments with tetrahedron with an A|C stack with 6 nucleotide sticky end.** The formed nanostructures were incubated at various temperatures for 1 hour as indicated in the lane. The intact fraction of tetrahedron was fitted with Boltzmann model to obtain the melting temperature for each experiment.

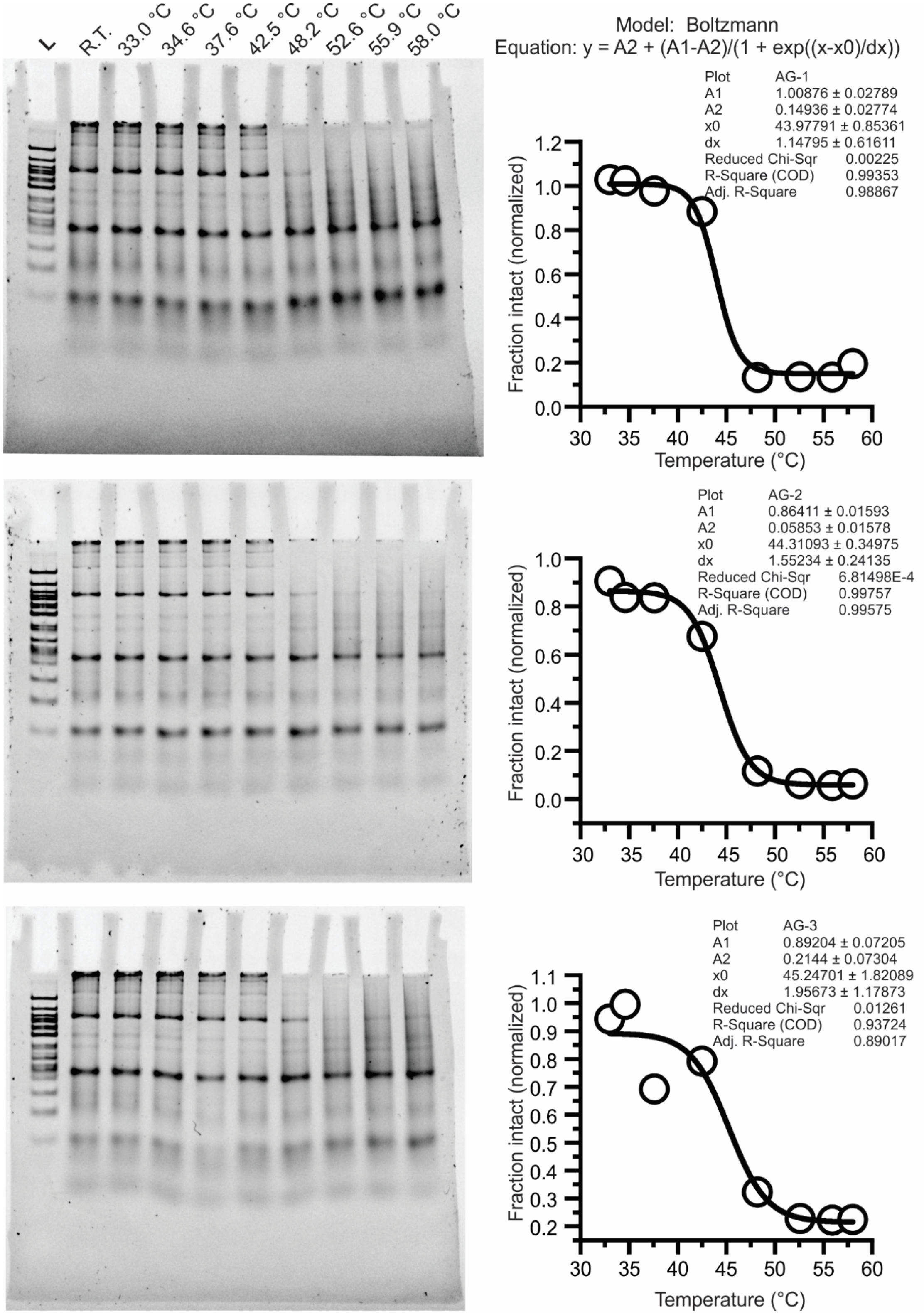


**Figure S5: Triplicate melting experiments with tetrahedron with an A|G stack with 6 nucleotide sticky end.** The formed nanostructures were incubated at various temperatures for 1 hour as indicated in the lane. The intact fraction of tetrahedron was fitted with Boltzmann model to obtain the melting temperature for each experiment.

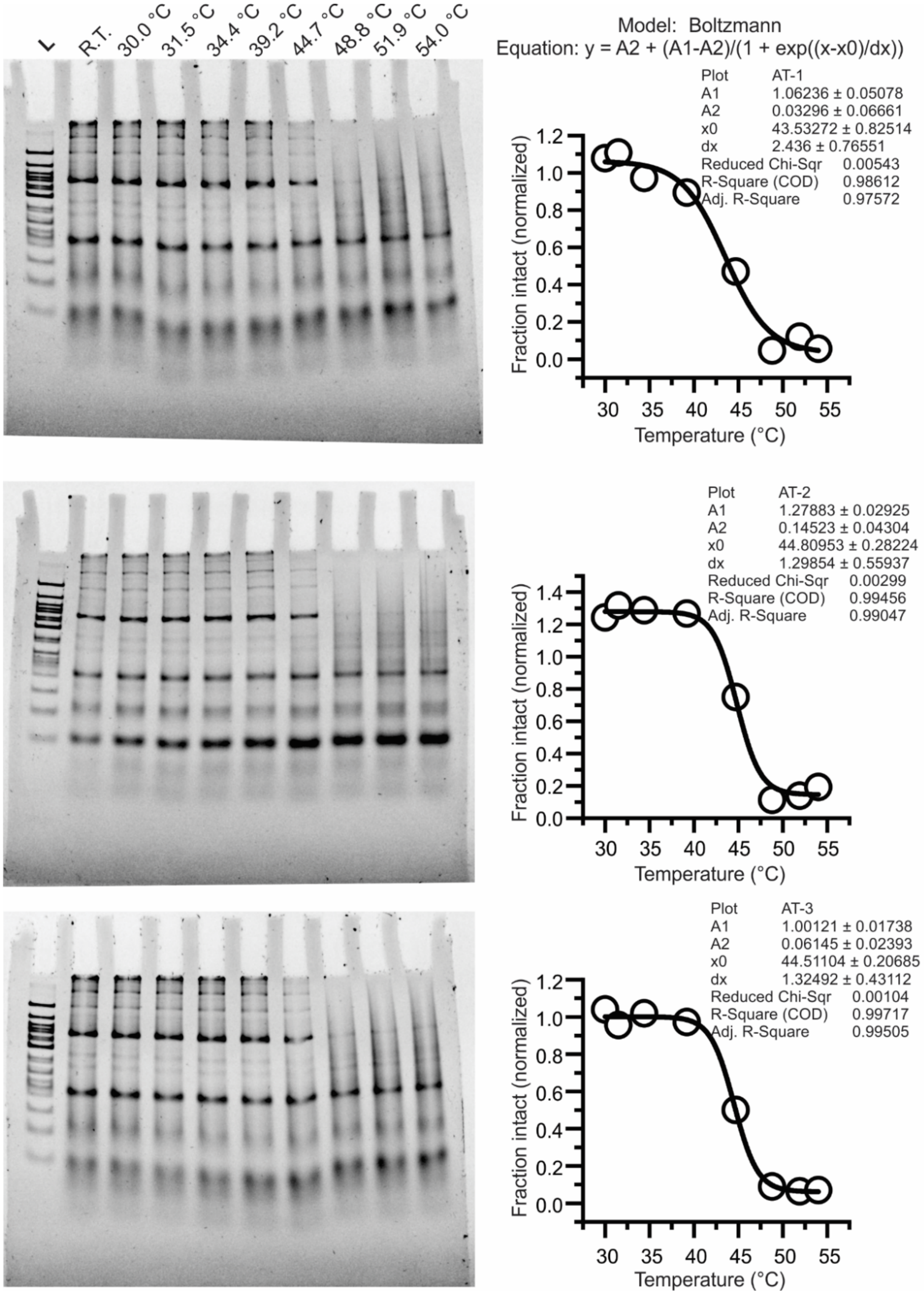


**Figure S6: Triplicate melting experiments with tetrahedron with an A|T stack with 6 nucleotide sticky end.** The formed nanostructures were incubated at various temperatures for 1 hour as indicated in the lane. The intact fraction of tetrahedron was fitted with Boltzmann model to obtain the melting temperature for each experiment.

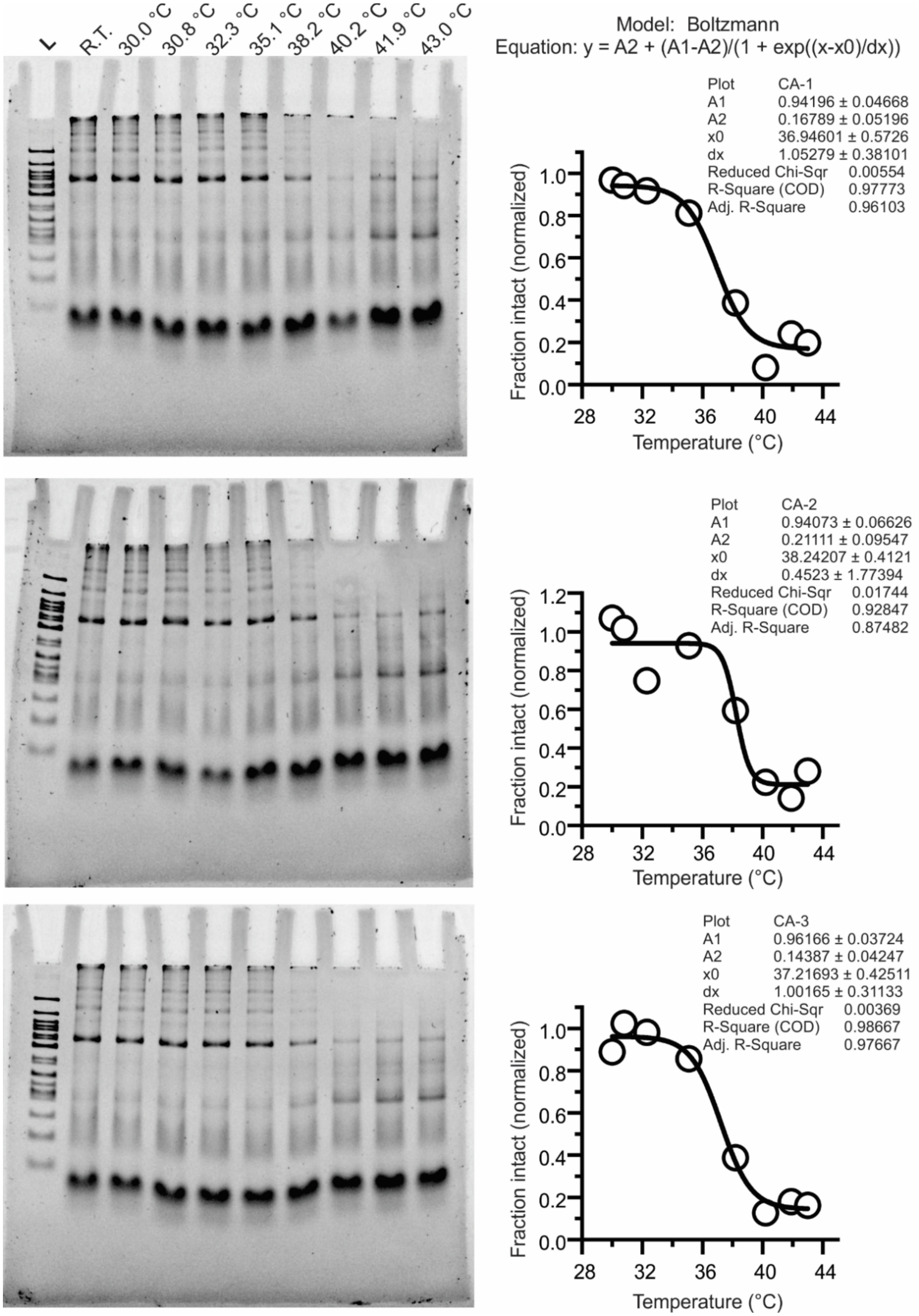


**Figure S7: Triplicate melting experiments with tetrahedron with a C|A stack with 6 nucleotide sticky end.** The formed nanostructures were incubated at various temperatures for 1 hour as indicated in the lane. The intact fraction of tetrahedron was fitted with Boltzmann model to obtain the melting temperature for each experiment.

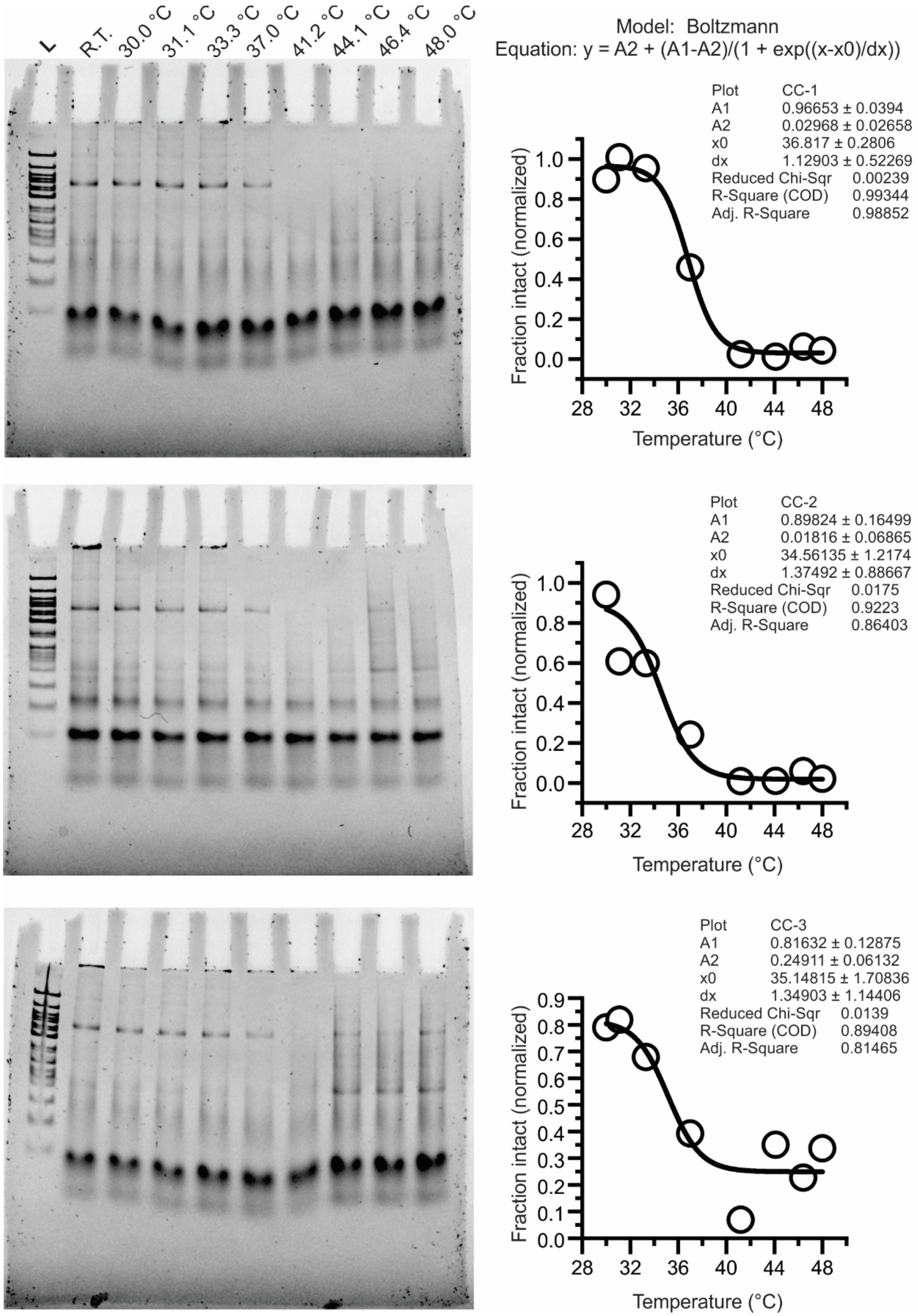


**Figure S8: Triplicate melting experiments with tetrahedron with a C|C stack with 6 nucleotide sticky end.** The formed nanostructures were incubated at various temperatures for 1 hour as indicated in the lane. The intact fraction of tetrahedron was fitted with Boltzmann model to obtain the melting temperature for each experiment.

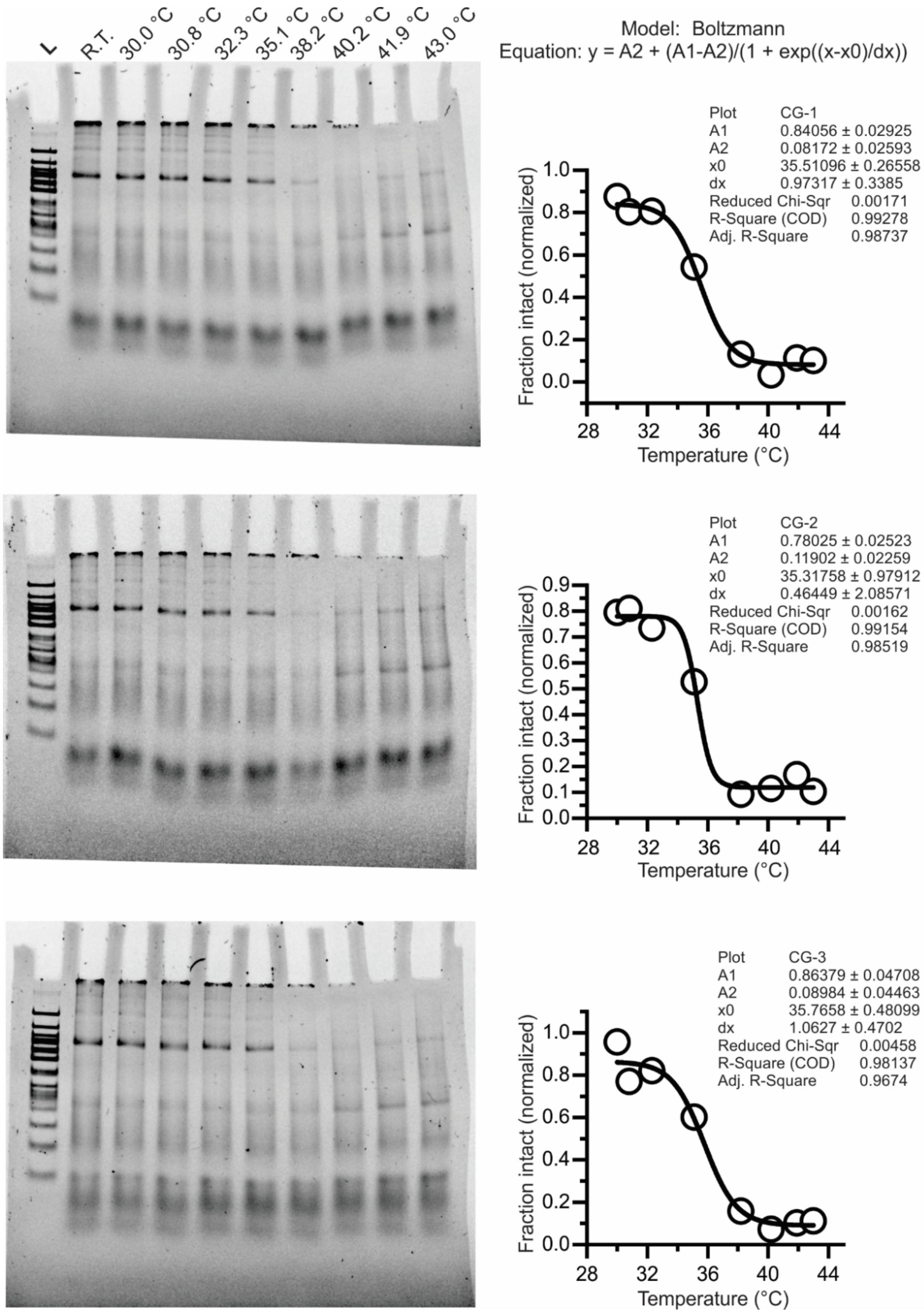


**Figure S9: Triplicate melting experiments with tetrahedron with a C|G stack with 6 nucleotide sticky end.** The formed nanostructures were incubated at various temperatures for 1 hour as indicated in the lane. The intact fraction of tetrahedron was fitted with Boltzmann model to obtain the melting temperature for each experiment.

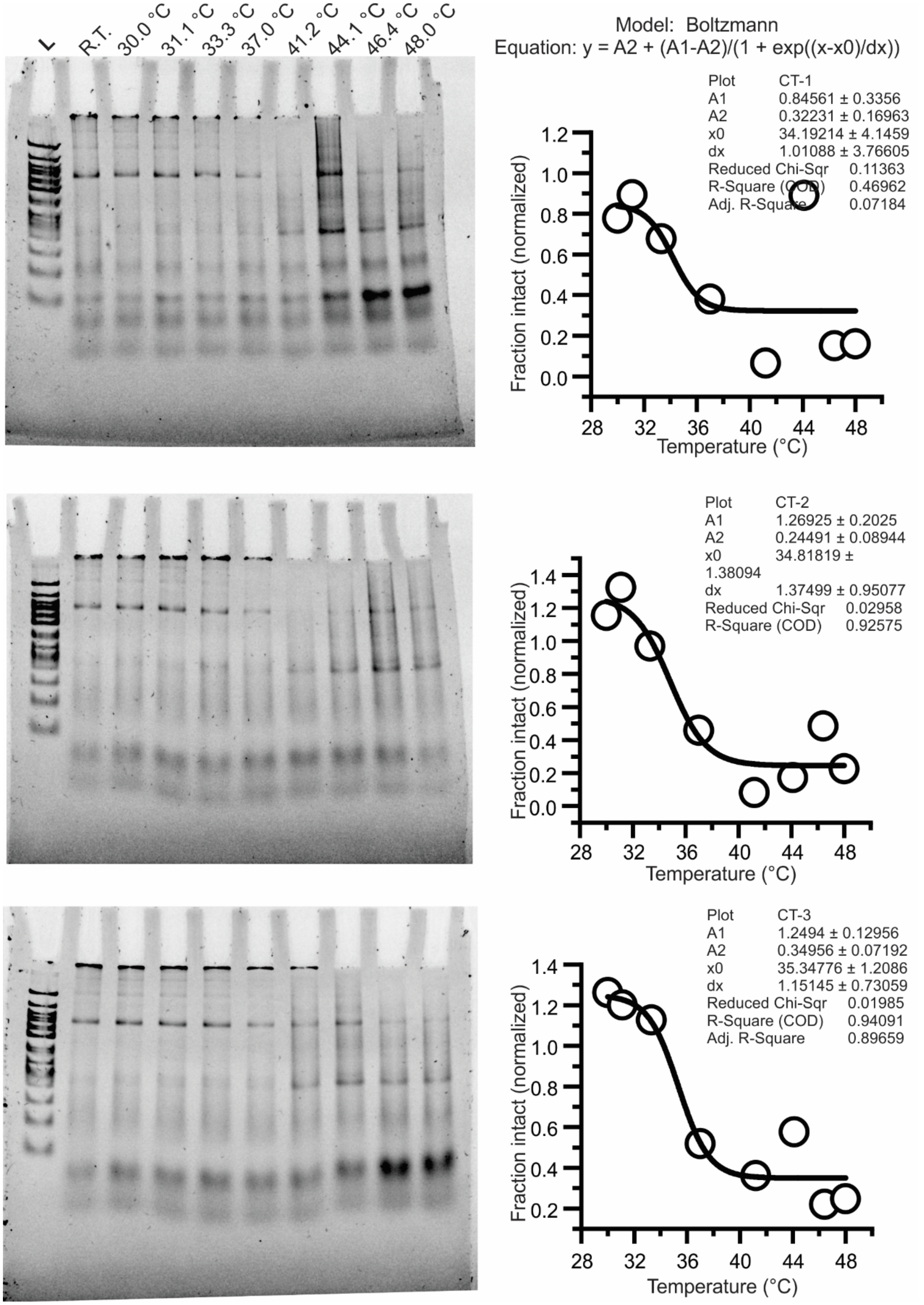


**Figure S10: Triplicate melting experiments with tetrahedron with a C|T stack with 6 nucleotide sticky end.** The formed nanostructures were incubated at various temperatures for 1 hour as indicated in the lane. The intact fraction of tetrahedron was fitted with Boltzmann model to obtain the melting temperature for each experiment.

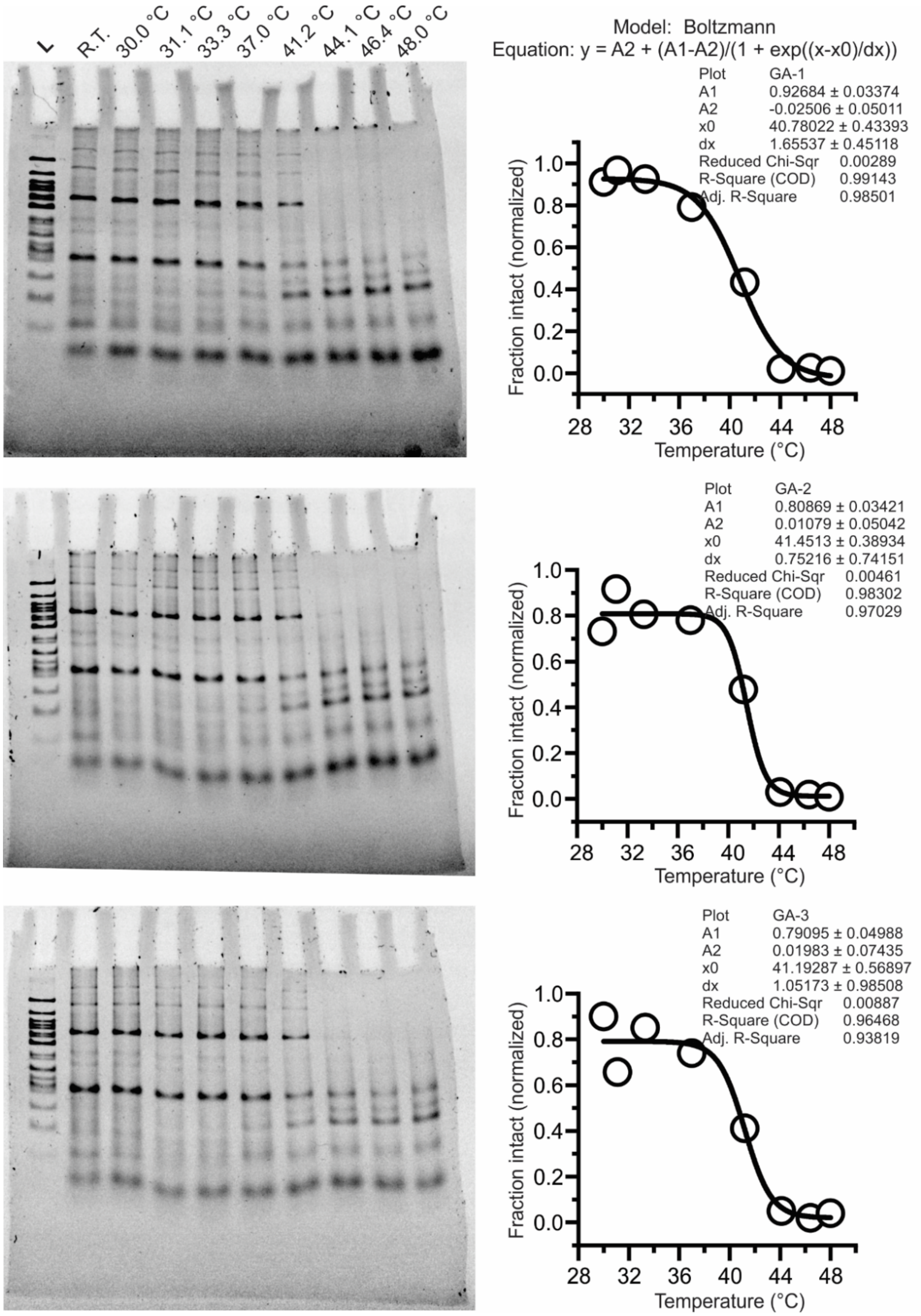


**Figure S11: Triplicate melting experiments with tetrahedron with a G|A stack with 6 nucleotide sticky end.** The formed nanostructures were incubated at various temperatures for 1 hour as indicated in the lane. The intact fraction of tetrahedron was fitted with Boltzmann model to obtain the melting temperature for each experiment.

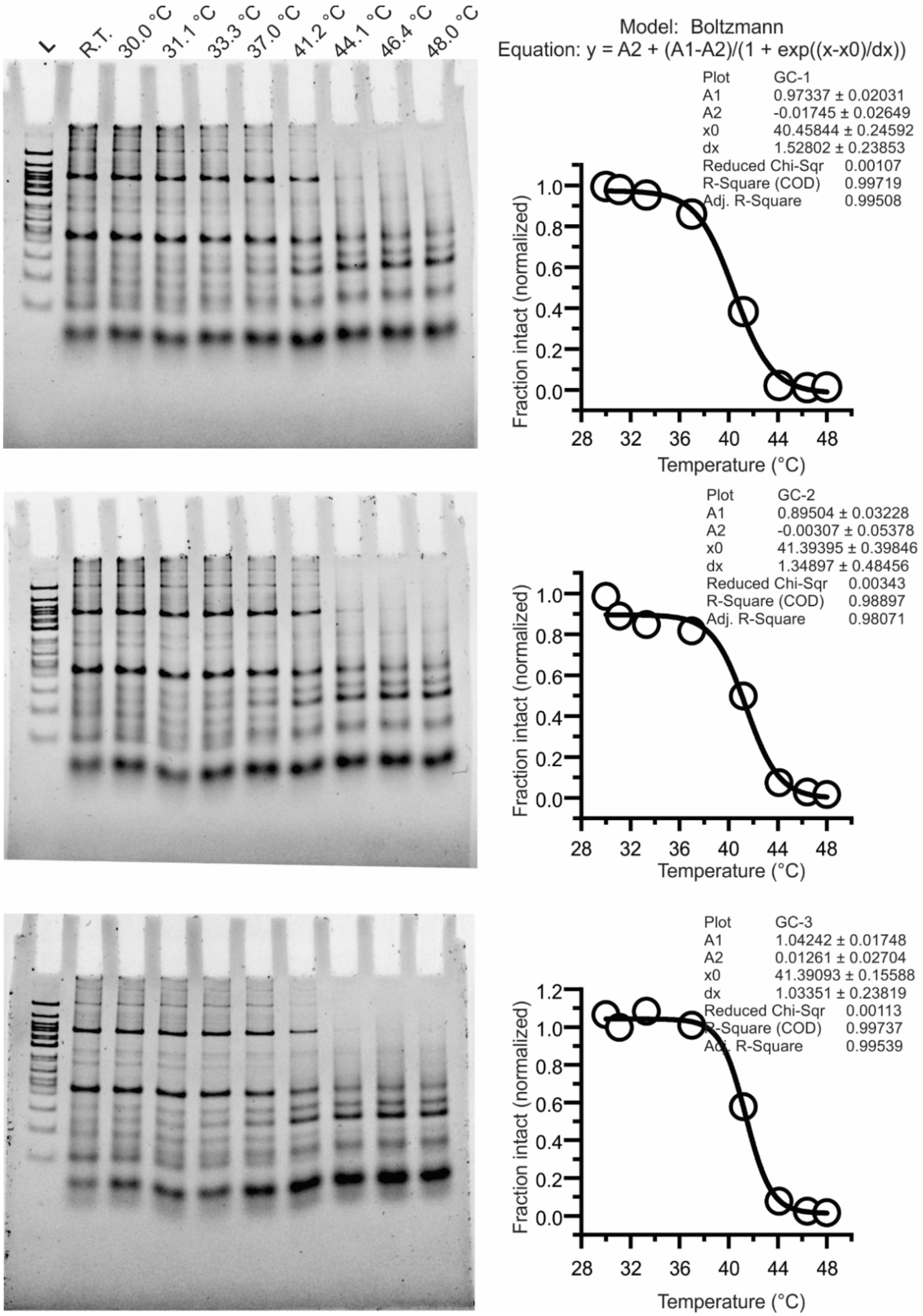


**Figure S12: Triplicate melting experiments with tetrahedron with a G|C stack with 6 nucleotide sticky end.** The formed nanostructures were incubated at various temperatures for 1 hour as indicated in the lane. The intact fraction of tetrahedron was fitted with Boltzmann model to obtain the melting temperature for each experiment.

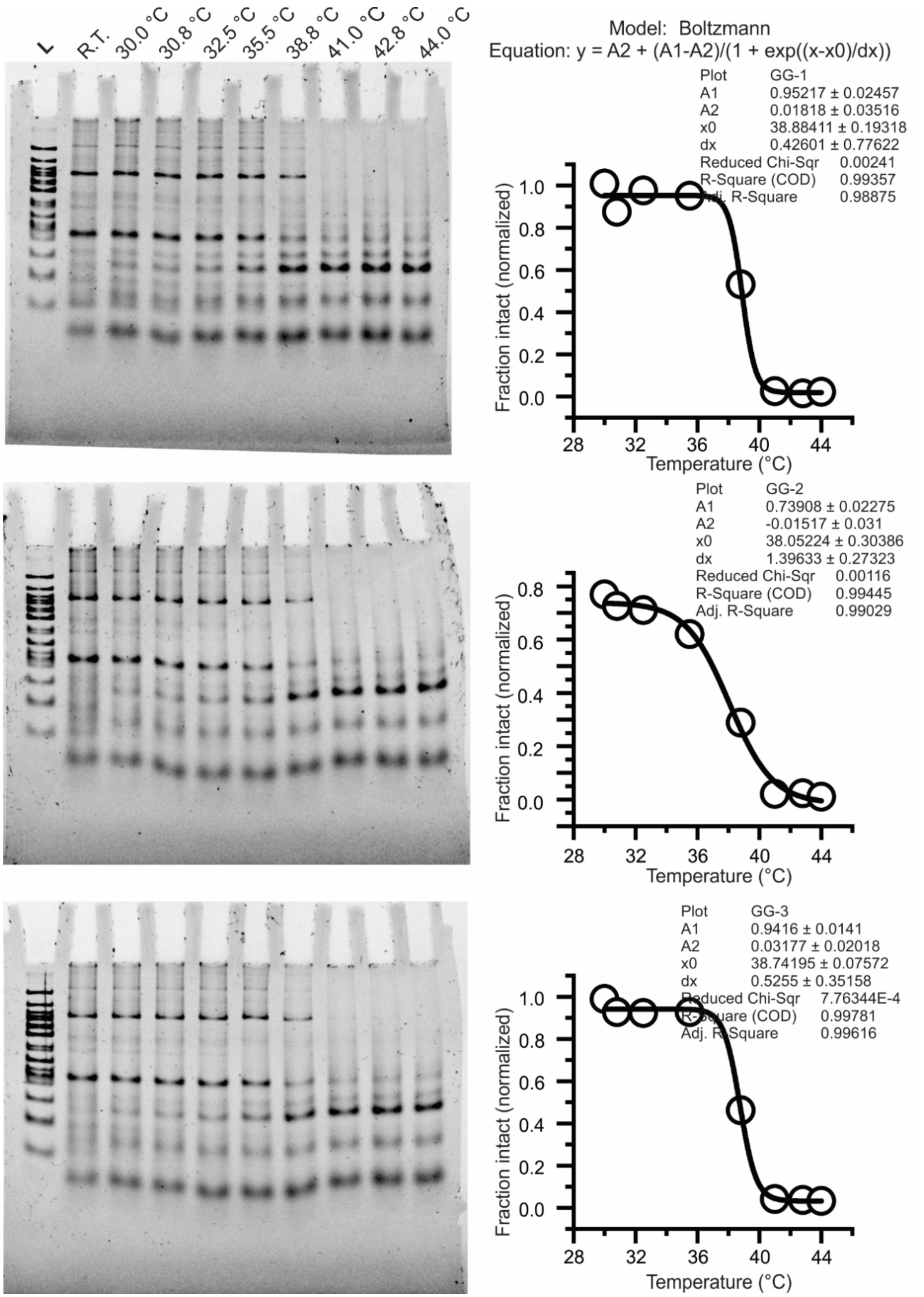


**Figure S13: Triplicate melting experiments with tetrahedron with a G|G stack with 6 nucleotide sticky end.** The formed nanostructures were incubated at various temperatures for 1 hour as indicated in the lane. The intact fraction of tetrahedron was fitted with Boltzmann model to obtain the melting temperature for each experiment.

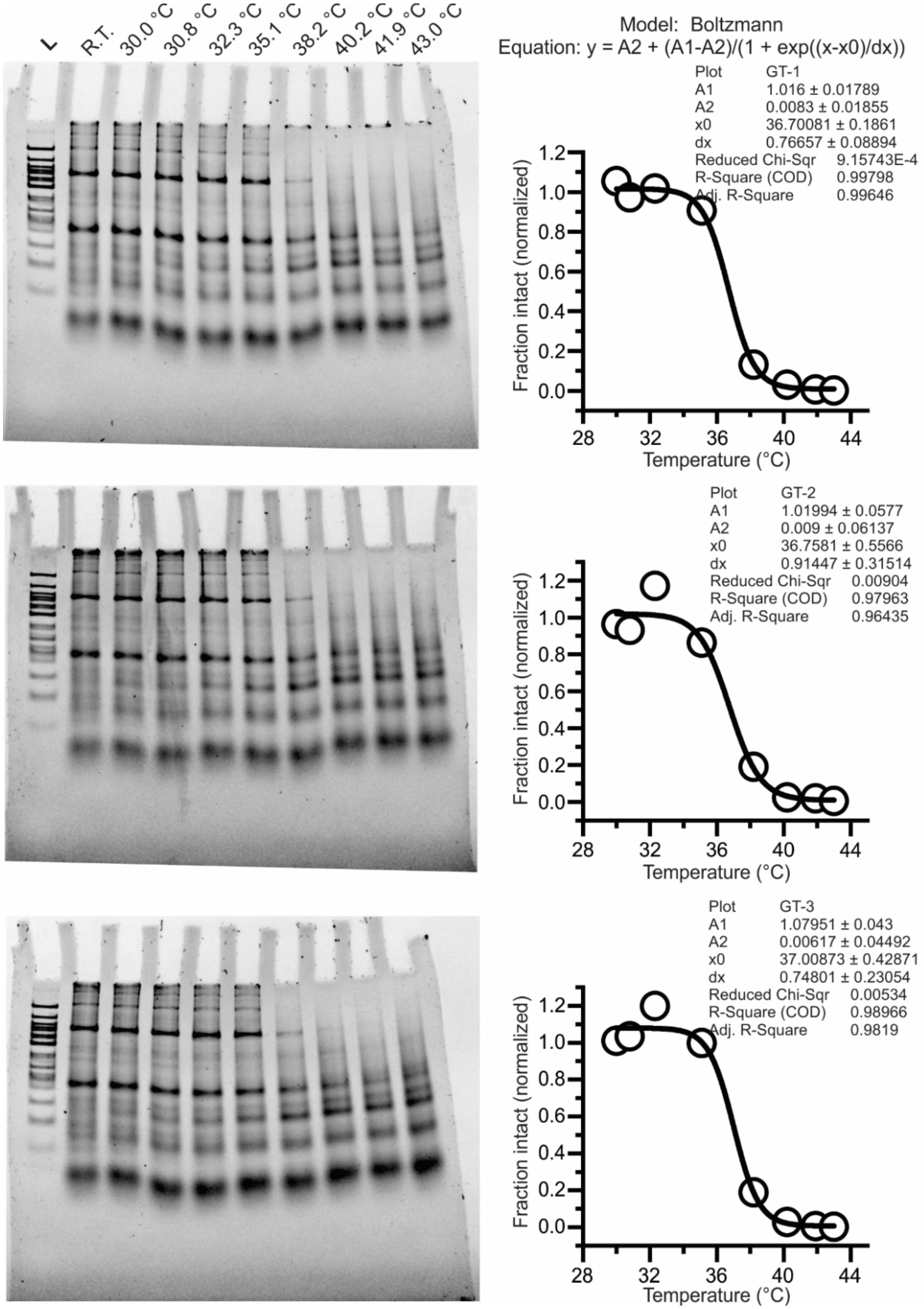


**Figure S14: Triplicate melting experiments with tetrahedron with a G|T stack with 6 nucleotide sticky end.** The formed nanostructures were incubated at various temperatures for 1 hour as indicated in the lane. The intact fraction of tetrahedron was fitted with Boltzmann model to obtain the melting temperature for each experiment.

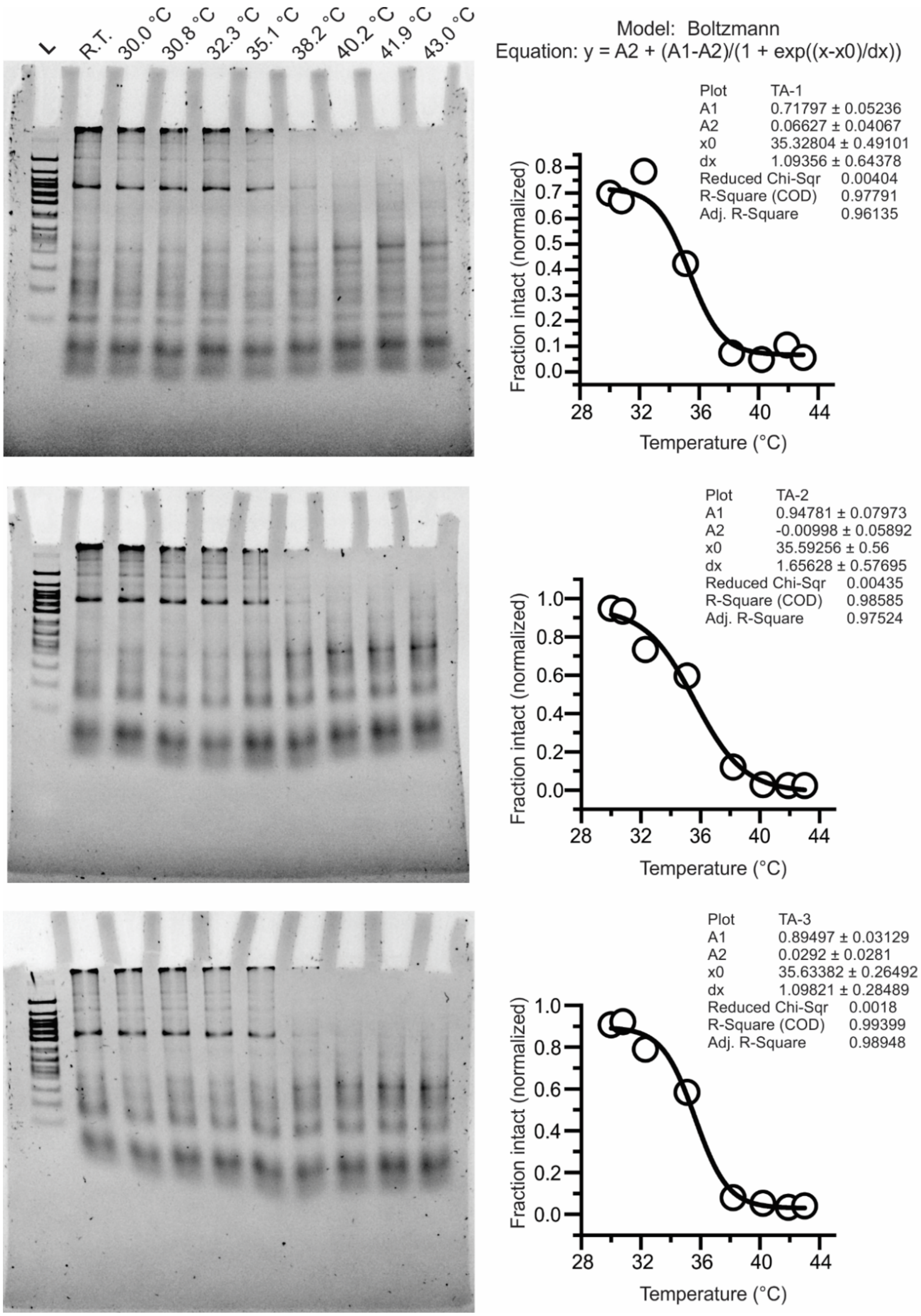


**Figure S15: Triplicate melting experiments with tetrahedron with a T|A stack with 6 nucleotide sticky end.** The formed nanostructures were incubated at various temperatures for 1 hour as indicated in the lane. The intact fraction of tetrahedron was fitted with Boltzmann model to obtain the melting temperature for each experiment.

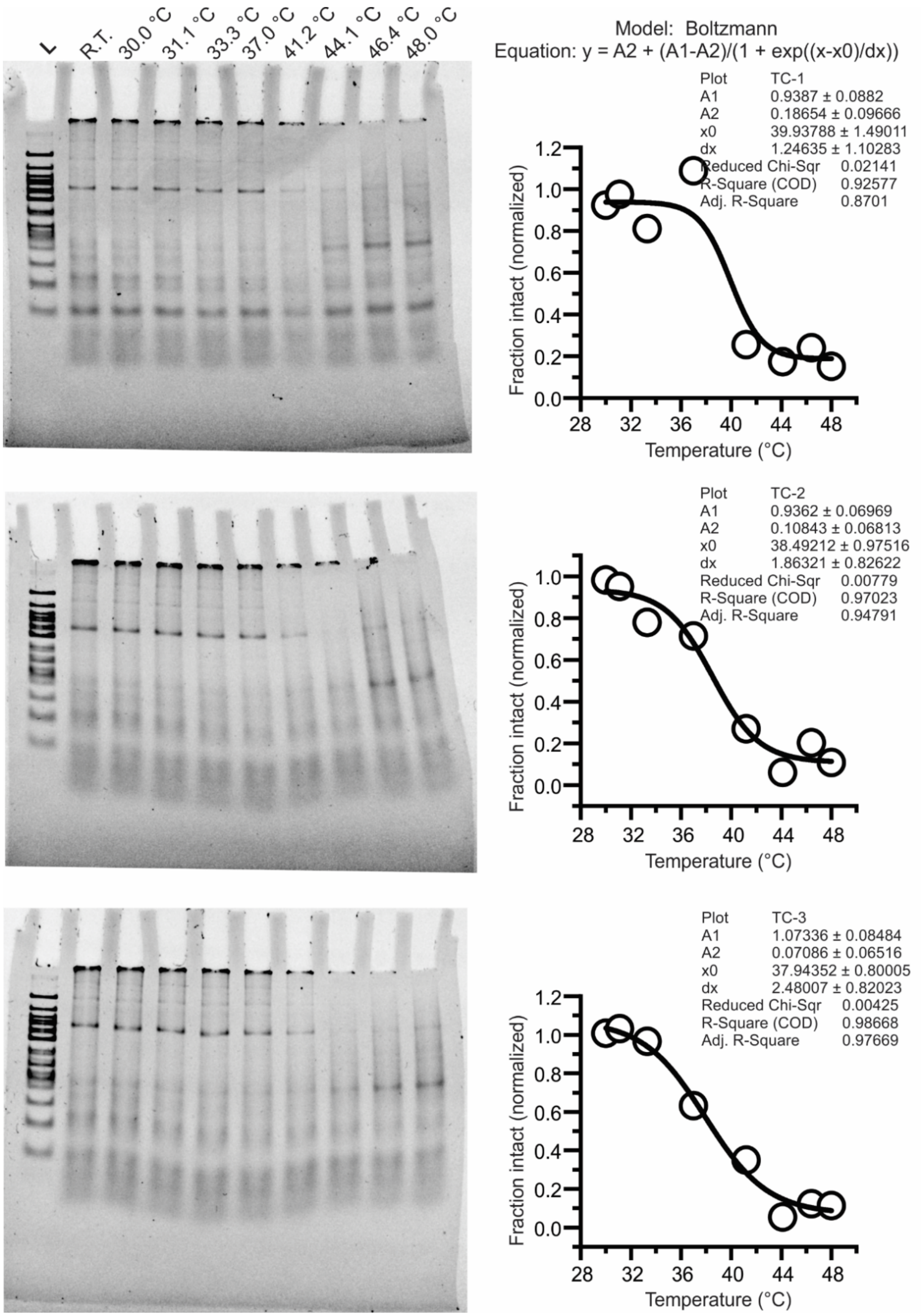


**Figure S16: Triplicate melting experiments with tetrahedron with a T|C stack with 6 nucleotide sticky end.** The formed nanostructures were incubated at various temperatures for 1 hour as indicated in the lane. The intact fraction of tetrahedron was fitted with Boltzmann model to obtain the melting temperature for each experiment.

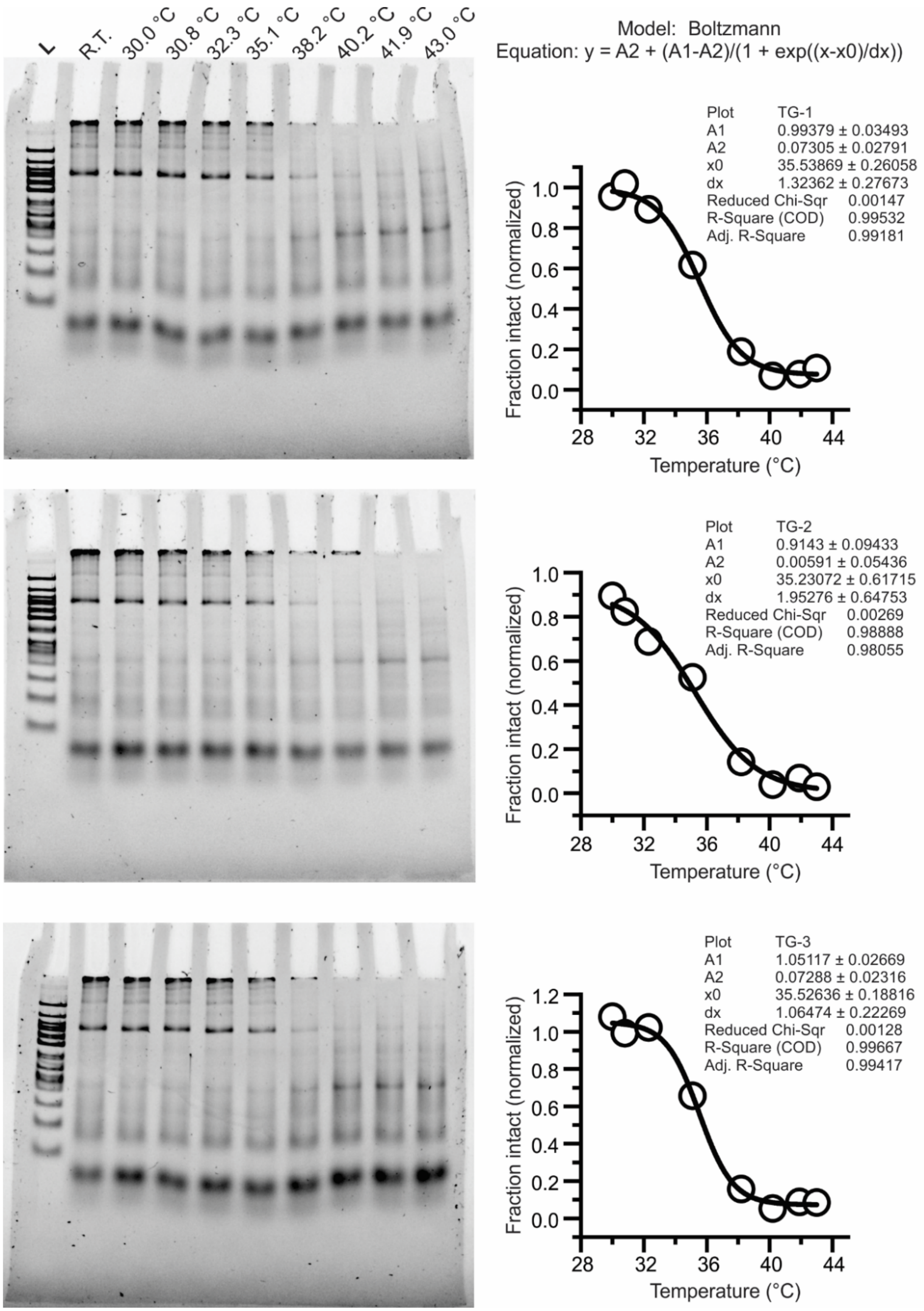


**Figure S17: Triplicate melting experiments with tetrahedron with a T|G stack with 6 nucleotide sticky end.** The formed nanostructures were incubated at various temperatures for 1 hour as indicated in the lane. The intact fraction of tetrahedron was fitted with Boltzmann model to obtain the melting temperature for each experiment.

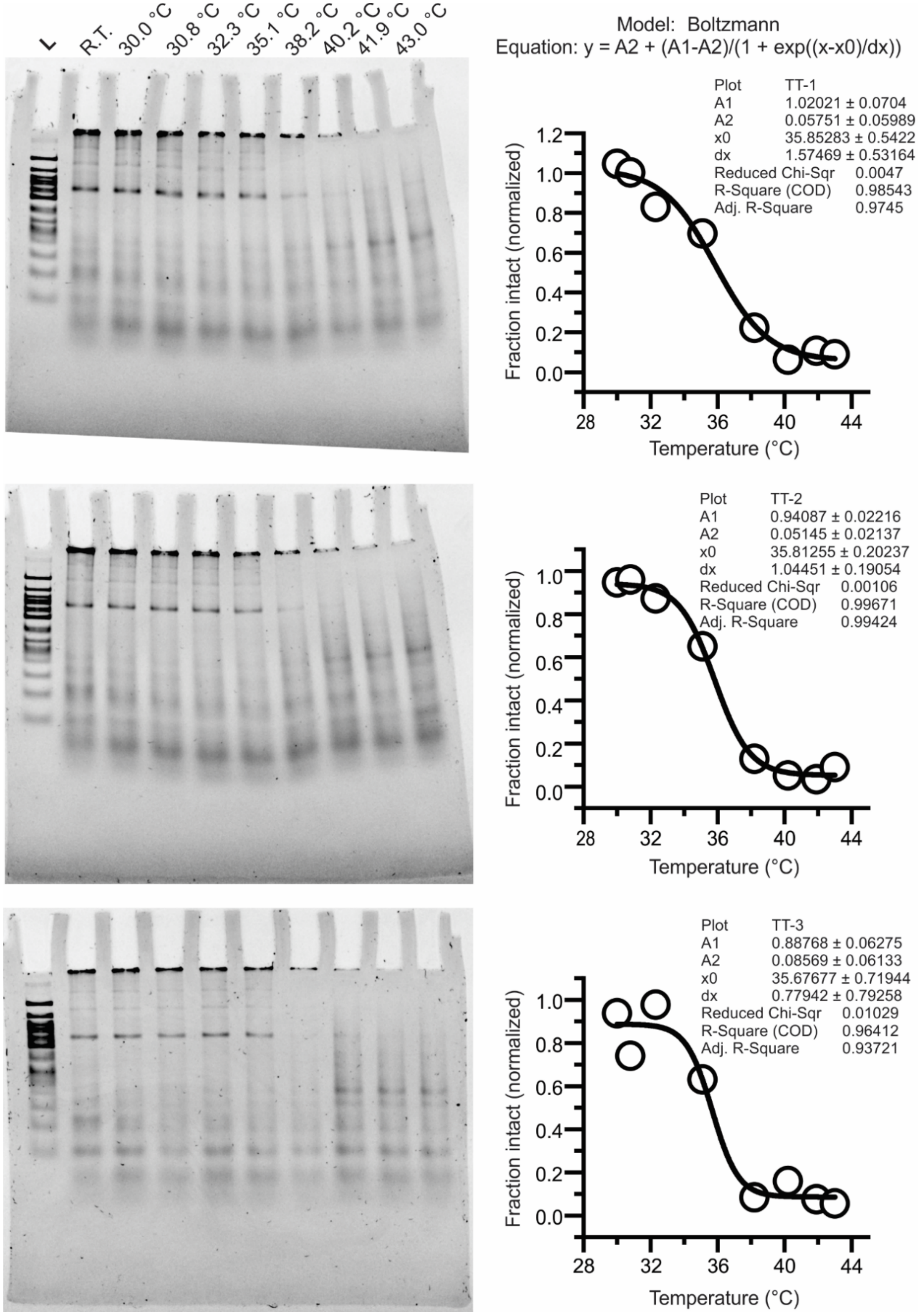


**Figure S18: Triplicate melting experiments with tetrahedron with a T|T stack with 6 nucleotide sticky end.** The formed nanostructures were incubated at various temperatures for 1 hour as indicated in the lane. The intact fraction of tetrahedron was fitted with Boltzmann model to obtain the melting temperature for each experiment.

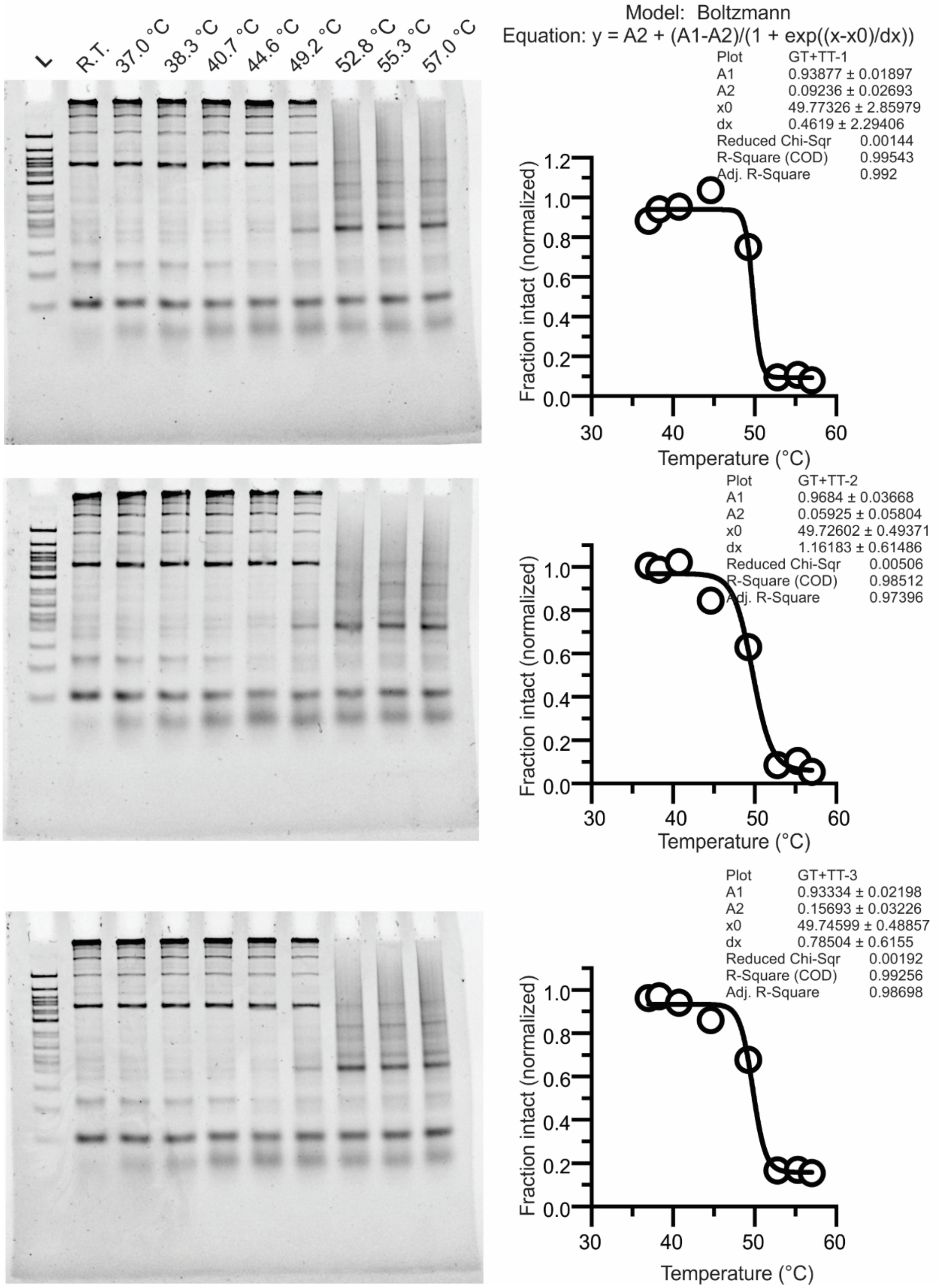


**Figure S19: Triplicate melting experiments with tetrahedron with a G|T and T|T stack with 6 nucleotide sticky end.** The formed nanostructures were incubated at various temperatures for 1 hour as indicated in the lane. The intact fraction of tetrahedron was fitted with Boltzmann model to obtain the melting temperature for each experiment.

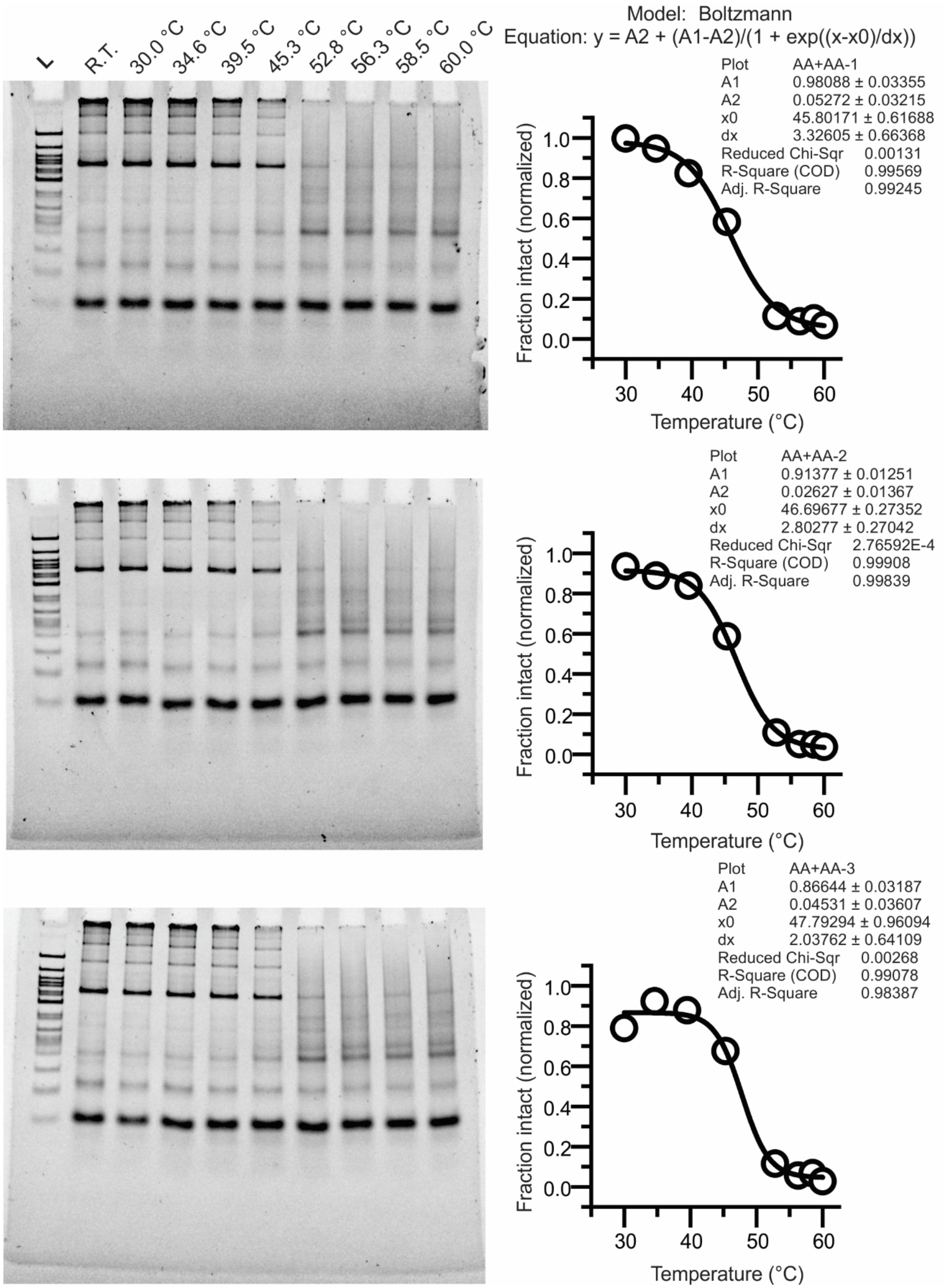


**Figure S20: Triplicate melting experiments with tetrahedron with 2 A|A stack and a 4 nucleotide sticky end.** The formed nanostructures were incubated at various temperatures for 1 hour as indicated in the lane. The intact fraction of tetrahedron was fitted with Boltzmann model to obtain the melting temperature for each experiment.

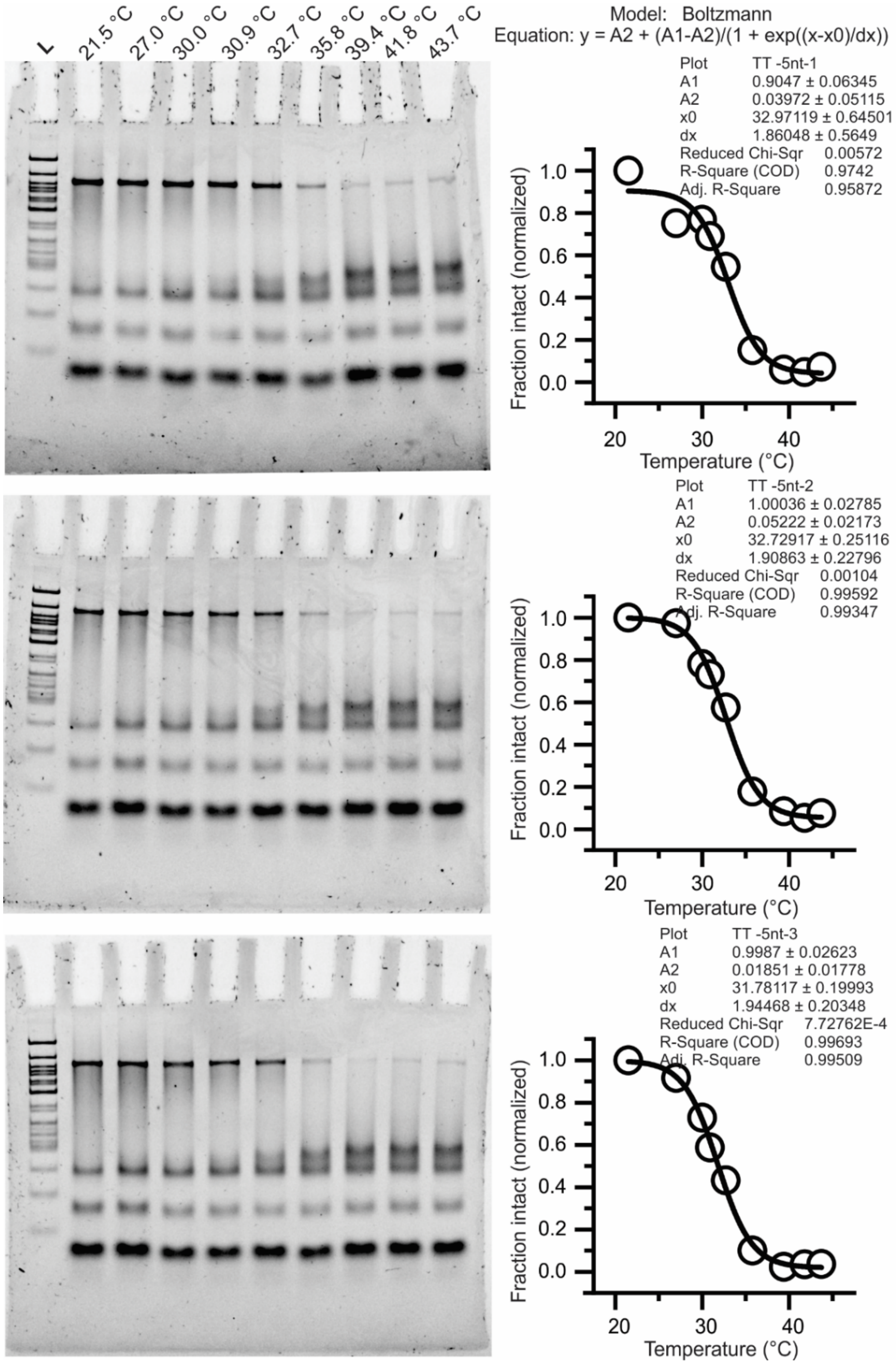


**Figure S21: Triplicate melting experiments with tetrahedron with 1 T|T stack and a 5 nucleotide sticky end.** The formed nanostructures were incubated at various temperatures for 1 hour as indicated in the lane. The intact fraction of tetrahedron was fitted with Boltzmann model to obtain the melting temperature for each experiment.

a

b

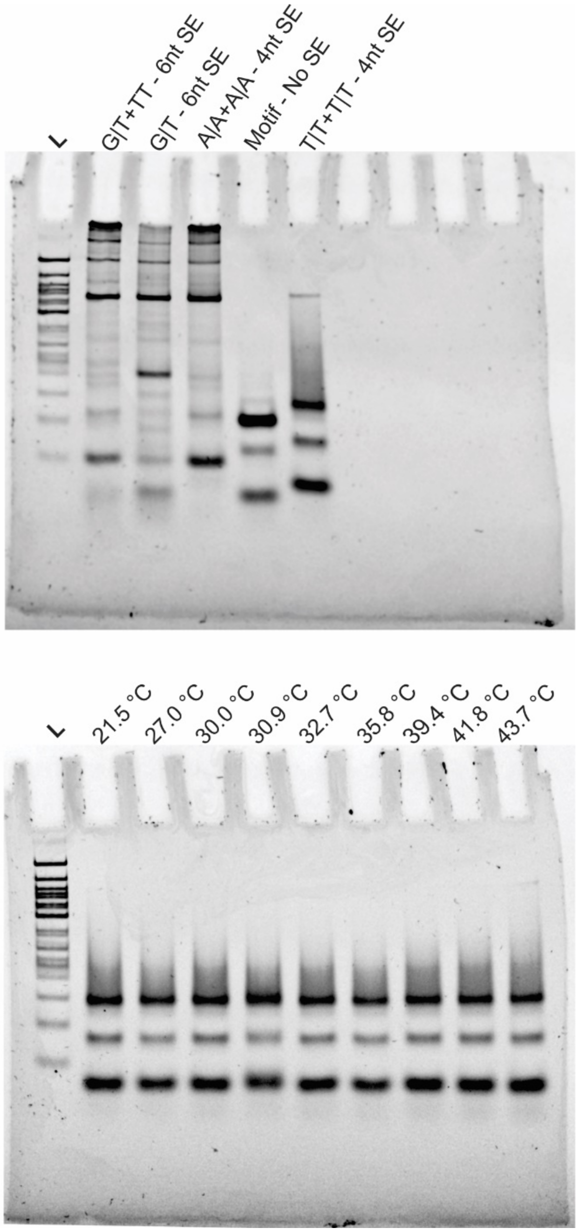


**Figure S22: Tetrahedron with 6 and 4 nt sticky ends with different base-stacks.** (a) PAGE confirmation formation of tetrahedra. Lane 1: 100 bp ladder, Lane 2: Tetrahedron with 6nt sticky end with G|T and T|T base stacks that are considered to be weak. Lane 3: Tetrahedron with 6nt sticky end with a single G|T stack. Lane 4: Tetrahedron with 4nt sticky end with 2 A|A stacks, which is the strongest stacks. Lane 5: DNA motif without sticky-end. Lane 6: Tetrahedron with 4nt sticky end with 2 T|T stacks, which is one of the weakest stacks and does not form reliably. (b) The tetrahedron with 4nt sticky ends and two T|T stacks was weak and not quantifiable during melting point determination.

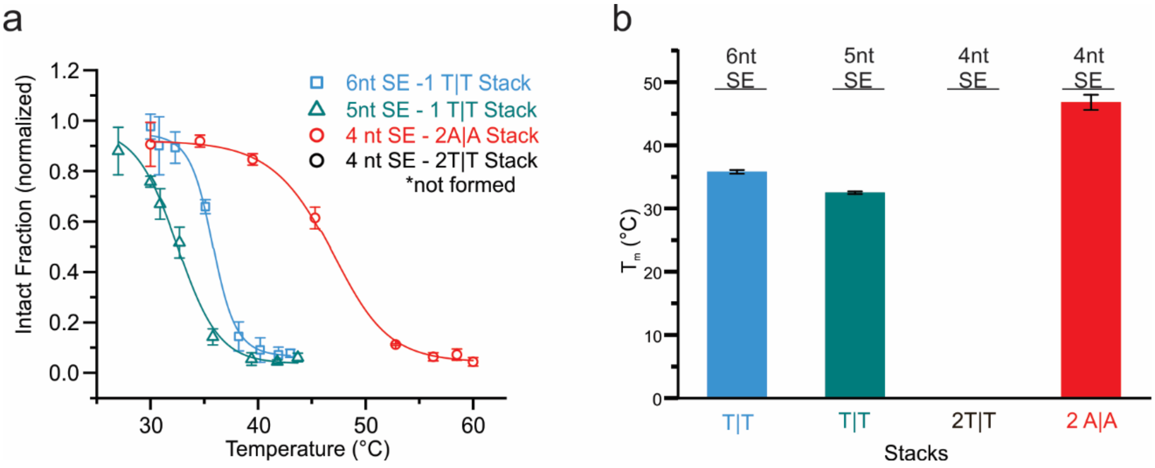


**Figure S23: Thermal stability difference between tetrahedron with T|T stacks and A|A stacks**. (a) The average melting profiles of the 3 tetrahedra that formed from triplicate measurements (Figures S18, S21, S22), fit with a Boltzmann curve to determine the melting temperatures. Tetrahedra with 4nt SE with 2 T|T stack failed to assemble (Figure S22), hence melting temperature wasn't determined. (b) A bar graphs representing the average meting temperatures of a 5 and 6 nt sticky end tetrahedron with a T|T stack and 4 nt sticky end tetrahedron with A|A stacks. The 4nt tetrahedron with 2 T|T stack failed to assemble. The error bars represent propagated error from the individual fits from Figure S18, S21 and S22. These experiments show that the reduced binding energy from shortened sticky-end can be compensated with stronger stacks.
